# Host specialization defines the emergence of new fungal plant pathogen populations

**DOI:** 10.1101/2024.09.30.615799

**Authors:** Wagner C. Fagundes, Rune Hansen, Idalia C. Rojas Barrera, Frauke Caliebe, Alice Feurtey, Janine Haueisen, Fatemeh Salimi, Alireza Alizadeh, Eva H. Stukenbrock

**Affiliations:** Environmental Genomics Group, Max Planck Institute for Evolutionary Biology, Plön & Christian-Albrechts University Kiel, Kiel, Germany; Laboratory of Evolutionary Genetics, Institute of Biology, University of Neuchâtel, Neuchâtel, Switzerland; Plant Pathology Group, Institute of Integrative Biology, ETH Zürich, Zürich, Switzerland; Department of Plant Protection, College of Agriculture and Natural Resources, Faculty of Agriculture, University of Tehran, Karaj, Iran; Department of Plant Protection, Faculty of Agriculture, Azarbaijan Shahid Madani University, Tabriz, Iran

## Abstract

Host-driven selection can be considered a strong driver of pathogen evolution. To successfully infect, colonize and complete their life cycle, plant pathogens are under constant selective pressures imposed by hosts, leading to genetic adaptation and possibly lineage radiation or speciation. Population and comparative genomics approaches are powerful tools to identify signatures of selection associated with host specialization in pathogen genomes and further allow recapturing population histories. Implementing such approaches, we identified evolutionary signatures of divergent host specialisation in distinct lineages of the fungal pathogen *Zymoseptoria tritici*, a major disease causing-agent of wheat. Unique collections of *Z. tritici* were isolated from wild (*Aegilops* spp.) and domesticated (*Triticum aestivum*) host grasses in the Middle East and whole-genome sequencing was performed in a selected subset of isolates from each collection. We observed distinct population structure between the two host-diverging pathogens and identified particular genomic features in the *Aegilops*-infecting isolates that may have shaped their evolutionary history. Phylogenomic analyses revealed that *A. cylindrica* and *A. tauchii* -infecting populations of *Z. tritici* form separate clusters, possibly reflecting incipient speciation driven by divergent host specialization. Using infection experiments, we confirm that *Z. tritici* isolates collected from *Aegilops* spp. only infect their respective host species and not *T. aestivum*. Population genomics analyses and demographic inference furthermore allowed us to detect signatures of recent selection and show that divergence of the wheat-infecting lineage likely coincided with wheat domestication. At last, we confirm a virulence-related role for one candidate effector located in a selective sweep region of the *A. cylindrica*-infecting pathogen. Taken together, our findings highlight the interplay between agricultural and wild hosts on the evolution of fungal plant pathogens and illustrate host specialization as a possible route of rapid pathogen emergence.

## Introduction

Host specialization is considered a strong driver of pathogen evolution [1–4]. Pathogen populations may diverge as they specialize to distinct hosts and ultimately become reproductively isolated lineages. In such cases, the different host species act as ecological barriers which prevent gene flow between differentially adapted pathogens. Host specialization furthermore involve the differentiation of intrinsic factors including physiological, molecular and biochemical properties that allow the invasion and persistence on specific hosts (e.g, virulence-related genes, antimicrobial metabolites, etc.) [1,5]. Depending on the intrinsic and extrinsic factors affecting the pathogen population on a specific host, reproductive isolation following host shift may be considered a driver of pathogen speciation and regarded a special case of ecological speciation [3,6,7].

Host adaptation is a continuous process in plant pathogens [8]. Plant immune receptors constantly evolve to recognize and overcome pathogen-produced virulence factors. The co-evolutionary process between pathogens and their hosts can be described as two distinct scenarios: a co-evolutionary “trench-warfare” or “arms race” scenario. The trench-warfare scenario describes the co-evolutionary dynamic as balancing or frequency-dependent selection, and the “arms-race” scenario as directional selection of favorable new alleles [9–12]. Typically, the genetic traits affected by trench-warfare and arms race evolution, are immune-receptors on the host side, and virulence-related genes, so-called effectors, on the pathogen side.

Effector proteins encoded by pathogen genomes are generally small, secreted proteins which very often lack sequence homology to other known proteins [13]. These small molecules can have very distinct functions, but are produced to interfere with host immune responses. Described effectors were shown to modulate host cell structures or interfere with plant metabolism through different mechanisms, often in a highly specific molecular dialogue [8,11,13–15]. Besides effectors, different classes of proteins (e.g. plant cell wall–degrading enzymes - PCWDEs and carbohydrate-degrading enzymes – CAZymes) and secondary metabolites (e.g. toxins) also play an important role in such host-pathogen interactions and can therefore also be under strong selective pressures imposed by plant hosts [13]. Footprints of strong positive selection have already been found in effector genes [16] as well as in toxins and genes encoding PCWDEs [17,18] in pathogen populations, suggesting the importance of such gene categories for pathogen fitness and adaptation to hosts.

Population genomics approaches provide a powerful resource to identify signatures of selection associated with speciation and host adaptation events besides recapturing the evolutionary history of species [19]. Adaptive changes and recent positive selection (e.g. selective sweeps), for example, can be identified through genomic scans as regions with skews in allele frequency distribution and distinct levels of linkage disequilibrium [20–24]. Selective sweep scans have previously been applied to fungal populations and allowed the identification of outlier regions containing genes potentially involved in host specialization and local adaptation [25–30].

*Zymoseptoria tritici* and closely related species have served as reference organisms for evolutionary genetic studies of plant-infecting fungal pathogens. *Zymoseptoria tritici* is a widely distributed wheat pathogen and the most devastating pathogen for wheat production in Europe [31]. The fungus causes the disease *Septoria tritici* blotch (STB) in wheat leaves, a disease that is characterized by a latent phase of biotrophy followed by a necrotic phase, in which host cell death and lesions become apparent. *Zymoseptoria tritici* reproduces both asexually and sexually and can undergo several cycles of reproduction in a year [32]. The levels of genetic variation and gene flow found in populations of *Z. tritici* around the world are extensive and levels of genetic variation within wheat fields are comparable to levels of variation found at global scales, indicating large effective population sizes and a high adaptive potential [33,34].

A study reconstructing the evolutionary history of *Z. tritici* indicated that it originated from a wild grass-associated ancestor in the fertile crescent region in the Middle East around 11,000 years ago, coinciding with the domestication of wheat and hence providing a great system to study recent pathogen emergence and host specialization [35,36]. Endemic relatives to *Z. tritici,* including *Z. pseudotritici* and *Z. ardabiliae* can be found in the Middle East as pathogens of different wild grass hosts [37,38]. Population genomics analyses of the “wild” *Z. pseudotritici* species revealed patterns of hybridization followed by a strong population bottleneck, leading to the emergence of a new hybrid species [39]. Moreover, recent studies suggest that recurrent introgression events between closely related species of *Zymoseptoria* occur at the center of origin in the Middle East and can leave footprints of highly variable regions in the genome [40,41]. However, the evolutionary mechanisms underlying such host-pathogen interactions in wild and agricultural ecosystems are still poorly understood.

In this study, we aimed to identify evolutionary and molecular signatures of host adaptation using host-divergent *Z. tritici* isolates as models of our study. Unique collections of *Z. tritici* strains were isolated from wild grasses belonging to the genus *Aegilops* and from cultivated common wheat (*Triticum aestivum*) in the Middle East. We used this resource to test the hypothesis that the different host species drive the evolution of new pathogen lineages in natural as well as agricultural habitats. Implementing population genomics analyses, we observed distinct population structure between the sympatric host-diverging populations and particular genomic features in the *Aegilops*-infecting *Z. tritici* isolates that may have shaped their evolutionary history. Genomic scans for selective sweeps identified diverse genomic regions under recent positive selection containing candidate genes potentially involved in host adaptation. Moreover, demographic inference suggests that divergence of the *Z. tritici* populations on wild and domesticated hosts likely occurred after wheat domestication. Using comparative infection experiments, we show that *Z. tritici* isolates collected from *Aegilops* spp. only infect *Aegilops* species and not *T. aestivum*, indicating a high degree of host specialization. At last, we confirmed a pathogenicity-related role for one of the three candidate effector genes present in the selective sweep region of the *Aegilops*-infecting *Z. tritici* population through a reverse genetics approach. Based on our findings, we propose that these effectors may be determinants of host specificity in this important class of plant pathogens.

## Results

### Isolation and characterization of *Z. tritici* isolates on wheat and wild wheat relatives

Wheat and *Aegilops* spp. samples presenting *Zymoseptoria* symptoms were collected at different locations in Iran in three consecutive years (2018-2020). From these samples, we isolated fungal specimens and confirmed the species using *ITS* and *Beta-tubulin* amplicon sequencing following standardized protocols and primers described previously [42]. A total of 310 fungal isolates were collected (135 from *Aegilops* spp. and 175 from wheat). Amplicon sequencing followed by nucleotide BLAST (blastn) analyses [43] revealed that all isolates collected from *Aegilops* spp. and wheat belong to the *Zymoseptoria tritici* species. Next, we performed clone correction through PCR using ISSR (Inter Simple-Sequence Repeat) markers to exclude highly similar isolates (e.g. clones) from our sequencing panel (see Methods). From the 310 initial isolates, we selected, based on the ISSR genotyping, a subset of 123 isolates for whole genome sequencing and subsequent analyses (48 isolated from *Aegilops* spp. and 75 isolated from wheat; S1 Table).

### Host-diverging *Z. tritici* isolates exhibit different levels of genomic diversity and linkage disequilibrium

Prior to assessing the population structure of the 123 sequenced *Z. tritici* isolates, we aimed to compare the overall genomic similarity and the levels of nucleotide diversity and linkage disequilibrium (LD) within each host-diverging collection from Iran. These analyses allowed us to have a first assessment of the overall genomic structure of the host-diverging isolates and to detect potential remaining clones in our sequencing dataset.

At first, we further corrected for clonality by assessing the genomic relatedness between the sequenced *Z. tritici* isolates. Hereby, we detected highly similar individuals (i.e. potential clones) by calculating genome-wide rates of Identity-By-State (IBS) among all 123 sequenced *Z. tritici* isolates. To this end, we used a quality-filtered dataset of 1,357,300 SNPs distributed among the thirteen core chromosomes of *Z. tritici* (see Methods). Based on a pairwise threshold of IBS > 0.9999, we excluded highly similar individuals from the dataset as they can represent clonal isolates. This filtering step resulted in a removal of 8 (out of 48) isolates and a reduction of 0.04% in the total number of SNPs for the Iranian *Aegilops*-infecting collection. For the Iranian wheat-infecting collection, we removed 30 (out of 75) isolates and observed a decrease of 0.12% in the total number of SNPs. From the 85 remaining isolates, we observed higher rates of IBS similarity within the *Aegilops*-infecting isolates (IBS similarity mean: 0.85) when compared to wheat-infecting ones (IBS similarity mean: 0.79) (Fig 1A and S1 Fig). This result suggests a difference in the overall genotypic composition between the two host-diverging Iranian collections, in which the *Aegilops*-infecting isolates show a higher genetic similarity when compared to the wheat-infecting ones.

**Fig 1.**
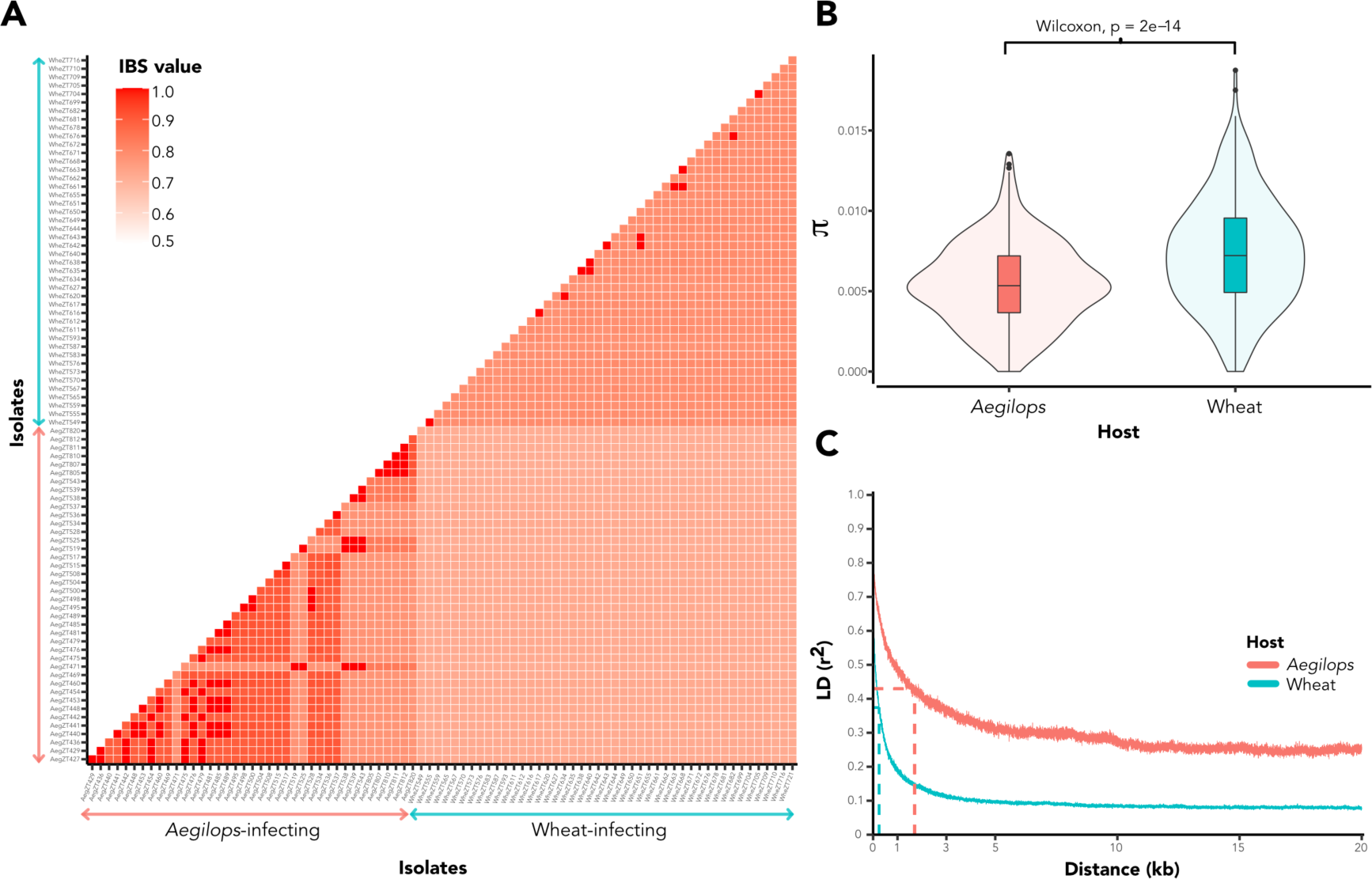
Host-diverging *Z. tritici* collections are genomically distinct. **(A)** Pairwise matrix representing genome-wide Identity-By-State (IBS) values between *Aegilops-* (red arrows) and wheat-infecting (blue arrows) *Z. tritici* isolates from Iran. Darker red colors represent higher IBS values and therefore higher similarity. **(B)** Distribution of genome-wide nucleotide diversity (*π*) values calculated for each host-diverging population individually. Values were calculated per 100 kb sliding windows. **(C)** Linkage disequilibrium decay across 20 kb of chromosome 1. Top, red line represents the LD decay for *Aegilops*-infecting isolates while the bottom, blue line represents the LD decay for the wheat-infecting population. Dashed perpendicular lines represent distance to reach 50% maximum LD value in each population.

Next, using the 85 isolate-filtered SNP dataset, we analyzed the levels of nucleotide diversity and assessed the linkage disequilibrium (LD) in each Iranian host-diverging population. For these analyses, we only used SNPs for each population with a MAF of ≥ 5%. This filtering step removed 25% and 32% of the total number of SNPs in the *Aegilops*- and wheat-infecting population, respectively. In line with the IBS results, we observed a lower genome-wide nucleotide diversity (*π*) for the *Aegilops*-infecting population (mean *π* = 0.005) when compared to the wheat-infecting population (mean *π* = 0.007) (Wilcoxon rank test, P-value = 2e-14, Fig 1B). We also observed a high differential LD decay between the two populations, in which *Aegilops*-infecting isolates had LD decayed to 50% of its maximum value at approximately 1580 bp and in the wheat-infecting population at approximately 230 bp (Fig 1C). Moreover, the *Aegilops*-infecting population exhibits a higher overall degree of LD between SNPs (mean LD r2 value: 0.30) than the wheat-infecting collection (mean LD r2 value: 0.10). Altogether, these results indicate that both *Z. tritici* collections are distinct at the genomic level and may have evolved under different demographic scenarios throughout history [44]. The differential high degree of LD and low nucleotide diversity in the *Aegilops*-infecting population further suggests that sexual reproduction is occurring less frequently among these isolates [45].

### *Aegilops*-infecting population form distinct *Z. tritici* clades

To determine patterns of relatedness among the *Z. tritici* populations from different hosts, we first estimated the phylogenetic relationship between the host-diverging *Z. tritici* populations using draft genome assemblies. First, we investigated the overall relatedness of *Z. tritici* populations using data from the Iranian populations as well as a population genomic dataset of wheat-infecting *Z. tritici* isolates collected across all major wheat growing regions of the world (S1 table). Neighbor-net network analysis based on whole genome distances revealed a clear separation between the *Aegilops*-infecting *Z. tritici* isolates and the ones isolated from wheat at different geographical locations (Fig 2A). We also observed, at finer scale, the segregation of two different *Aegilops*-infecting populations (Fig 2A). Similar results were obtained using a maximum-likelihood phylogenetic approach based on intraspecific genomic variation (PoMo; [46,47]) in which we also observed a clear separation between the *Aegilops*-infecting *Z. tritici* isolates from the wheat-infecting ones (S2 Fig). Genome-wide divergence values based on the Weir & Cockerham’s Fst fixation indices [48] also indicated that the two *Aegilops*-infecting populations were highly divergent from other *Z. tritici* populations as well as between each other (S3 Fig), suggesting that host specificity rather than geographical origin is determining *Z. tritici* population structure and population divergence.

**Fig 2.**
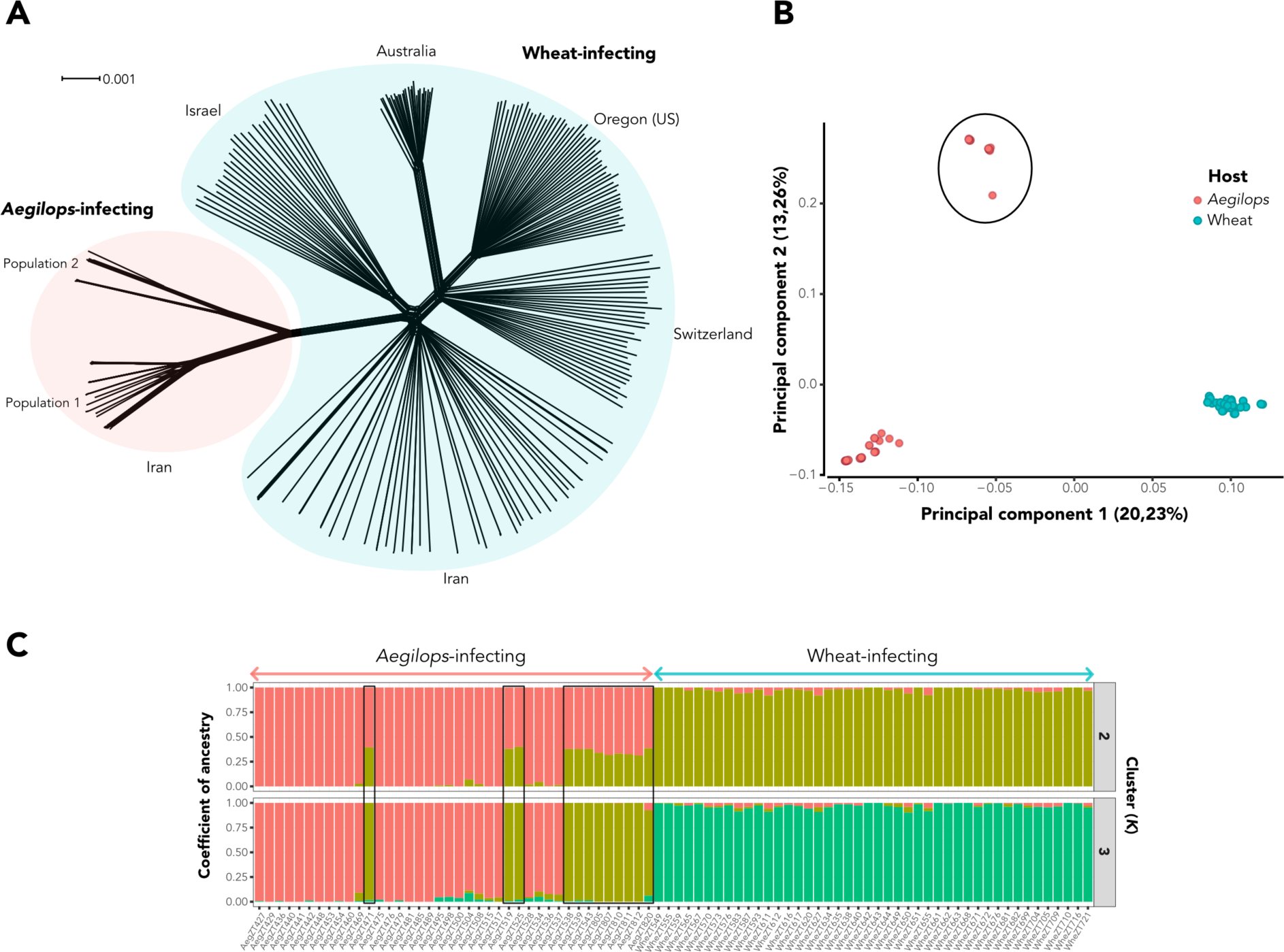
Population structure of Iranian *Z. tritici* isolates is correlated with host species. **(A)** Neighbor-net network based on whole genome distances. Red ellipse represents the two *Aegilops*-infecting *Z. tritici* populations while the blue elliptical shape represents worldwide wheat-infecting *Z. tritici* populations. Geographical locations of origin are also indicated. Scale bar indicates branch distances. **(B)** PCA analyses based on genome-wide independent SNPs. The first two principal components (PCs) are shown with the variance explained by each component in parenthesis. Red dots represent *Z. tritici* isolates collected from *Aegilops* species while blue dots represent the ones isolated from wheat. *Aegilops* population 2 is indicated with a black ellipse. **(C)** Ancestry coefficients and clustering assignments using the sNMF software. Each vertical bar represents an isolate from the *Aegilops* (red arrow) and wheat-infecting (blue arrow) *Z. tritici* collection with each color indicating one genetic cluster. The color height in each vertical bar represents the probabilities of cluster assignment based on genome-wide independent SNPs. Isolates belonging to the *Aegilops* population 2 are highlighted in black.

Next, considering the divergence between the host-diverging *Z. tritici* isolates, we asked whether the *Aegilops*-infecting populations were more closely related to other *Zymoseptoria* species, including species co-occurring in natural grassland vegetations in Iran, than to the wheat-infecting *Z. tritici*. To test this hypothesis, we used the same Neighbor-net network approach based on genomic distances including draft genome assemblies from the closely related *Zymoseptoria* species *Z. passerini*, *Z. ardabiliae* and *Z. brevis* (S1 Table) and observed an evident separation of the *Z. tritici* cluster from the closely related species (S4 Fig). We note however that several edges (internal branches connecting taxa nodes) connected the *Aegilops*-infecting isolates to the other *Zymoseptoria* species (S4 Fig). Phylogenetic network analyses can illustrate possible evolutionary relationships as hybridization and recombination between taxa, in which the edges represent such reticulate evolutionary events [49]. Based on these results, we suggest that the *Aegilops*-infecting isolates represent *Z. tritici* lineages distinct from the wheat-infecting one and further hypothesize that these lineages may have experienced gene flow events with other *Zymoseptoria* species throughout evolution.

### Population structure is correlated with host species

We assessed the population structure among host-diverging *Z. tritici* isolates using different methods. We hereby focused on *Z. tritici* isolates collected in the same geographical region in the Middle East and addressed the extent of gene flow between the pathogen populations with distinct hosts, but overlapping dispersal ranges. To this end, we only included the population genomic dataset of the *Aegilops*-infecting populations and the wheat-infecting population derived from agricultural field sites in Iran. We also restricted our population structure analyses to 2507 independent (i.e. un-linked) SNPs distributed across the thirteen core chromosomes of *Z. tritici*.

Principal component analysis (PCA) revealed an evident separation between the isolates collected from *Aegilops* spp. and from wheat (Fig 2B and S5 Fig). In line with the phylogenomic results, the *Aegilops*-infecting cluster also showed a further separation, in which a smaller subcluster was composed of twelve isolates obtained from *Aegilops* spp. Samples collected at a unique geographical location in two different years (Fig 2B). Corroborating these findings, clustering analysis based on ancestry coefficients implemented in the sNMF software [50,51] also showed a clear separation between the *Aegilops*- and wheat-infecting isolates and the further substructure within the *Aegilops*-infecting population (*K*=2 and *K*=3; Fig 2C). No large clustering overlaps were observed between the host-diverging populations at higher clustering models (*K*=4 and *K*=5; S6 Fig). Taking these results together, we could define three distinct Iranian *Z. tritici* populations. Hereafter, we refer to these populations as “*Aegilops* population 1”, representing the larger *Aegilops*-infecting *Z. tritici* population; “*Aegilops* population 2”, representing the smaller population of twelve *Aegilops*-infecting *Z. tritici* isolates and the “wheat population”, representing the wheat-infecting *Z. tritici* isolates.

Considering the population segregation observed for the *Aegilops* populations 1 and 2, we hypothesized that host specificity to different *Aegilops* species or genotypes could explain the observed population structure of the pathogen. Our initial characterization of the *Aegilops* material was only based on manual inspection of the herbarium plant material. We therefore further characterized the host leaves using PCR amplification of the *ITS2* barcode locus. Intriguingly, the *ITS2* sequences of leaf samples from the *Aegilops*-infecting population 1 revealed that the *Z. tritici* strains were isolated from *Aegilops cylindrica* or a closely-related *Aegilops* species (e.g. *A. triuncialis* or *A. tauschii*) while *Z. tritici* isolates from the *Aegilops* population 2 were obtained from *A. tauschii* or other closely-related *Aegilops* species (e.g. *A. geniculata*; S7 Fig and S2 Table). For the wheat-infecting *Z. tritici* isolates, amplicon analyses suggested that their hosts were indeed from the common wheat species (*Triticum aestivum* L.) or from closely-related polyploid wheat species (e.g. *T. durum*) (S7 Fig and S2 Table). Taken together, these results support the hypothesis that distinct host species are shaping the structure of the *Z. tritici* populations among closely related plant species in natural grassland vegetations.

### Mating type distribution differs between host-diverging populations

The high levels of linkage disequilibrium and IBS and low genome-wide nucleotide diversity observed in the *Aegilops*-infecting collection (Fig 1) led us to hypothesize that asexual propagation dominates in these *Z. tritici* isolates. One of the strongest pieces of evidence for clonal reproduction in ascomycete fungi with a heterothallic mating type system is an uneven distribution of mating type alleles in a population or the complete lack of one of the idiomorphs. Several genes are involved in mating, but two major idiomorphs (MAT1-1 and MAT1-2) are described in *Z. tritici* [52]. To analyze the distribution of these idiomorphs, we performed BLAST searches using nucleotide sequences of both mating type loci against whole-genome assemblies of the 123 sequenced individuals. We then counted the number of idiomorphs for each *Z. tritici* population individually (wheat population, *Aegilops* population 1 and the *Aegilops* population 2). Both the wheat and the *Aegilops* population 1 did not deviate from the even (1:1) frequency of mating types (*χ*2 P=0.204, wheat population and *χ*2 P=0.3173, *Aegilops* population 1), suggesting that sexual reproduction is occurring in these populations (S3 Table). In contrast, in the *Aegilops* population 2 no isolate had the MAT1-1 idiomorph suggesting a dominant role of asexual propagation (S3 Table). This is in agreement with the lack of LD decay and the low genome-wide nucleotide diversity that we observe in the *Aegilops* population 2 (S8 Fig). We note that our collection of *Aegilops* population 2 only includes twelve isolates collected at one location in Iran (Fig 2 and S1 Table). It is therefore possible that sampling in this particular location had an impact on the frequency of mating types and levels of diversity detected.

### *Aegilops*-infecting *Z. tritici* lineage diverged after wheat domestication around 5 Kya

The phylogenetic and population divergence observed between the *Z. tritici* populations infecting *Aegilops* spp. from the ones infecting wheat prompted us to further investigate the demographic history of these isolates. To this end, we used a Multiple Sequentially Markovian Coalescent MSMC2 method [53,54] to estimate the coalescence rates and divergence times between haplotypes from the wheat population and the *Aegilops* population 1. This method assumes propagation of generations by sexual recombination and we could therefore not perform the demography analyses with the *Aegilops* population 2.

To convert estimates of coalescence times, we assumed a mutation rate of 3.3e-^8^ per cell cycle as in estimated for *S. cerevisiae* [55] and assuming 1 generation of sexual reproduction per year, we find that the wheat-infecting population experienced a population expansion followed by a bottleneck between 30,000 and 10,000 years ago (S9A Fig). A similar pattern was identified in the *Aegilops* population 1, although we observe a large variation in effective population size (Ne) for more recent times (S9B Fig).

Next, to assess the timing of divergence between the two populations, we inferred the relative cross-coalescence rate (RCCR) based on eight haploid genomes (four from each population). The results suggested that divergence of the two *Z. tritici* populations co-occurred with wheat domestication around 10,000 years ago; reflected by a decrease of the RCCR at this time (S10 Fig). The average RCCR falls below 0.5 around at a time that corresponds to 4.415 Kyears (Standard Deviation = 1.617 Kyears). We found similar results when we permuted the samples between the two populations, indicating that this divergence signal is strong and well supported by the data (S10 Fig). Altogether, these findings suggest that the two host-diverging *Z. tritici* populations have diverged and expanded in a time frame from 5,000-10,000 years ago and corroborate our hypothesis that these populations represent distinct host-specific *Z. tritici* lineages. Moreover, the time frame of divergence detected here correlates with previous coalescence analyzes of the wheat-infecting *Z. tritici* population, which indicates that *Z. tritici* has split from its closest sister species *Z. pseudotritici* approximately 10,500 years ago coinciding with the domestication of its host wheat [36]. Regarding the isolates from the *Aegilops* population 2, additional sampling and demographic analyses are necessary to explore the potential divergence time of this population.

### *Aegilops*-infecting *Z. tritici* isolates are host specific

Considering the evolutionary divergence observed between the sympatric populations of *Z. tritici* infecting wheat and *Aegilops*, we investigated if the *Z. tritici* isolates from these populations also had distinct host ranges. To this end, we performed independent qualitative and quantitative *in planta* virulence assays in the greenhouse. In a first experiment, we inoculated three batches of sequenced isolates based on the identified populations, namely a “*Aegilops* pop 1” batch, and a “*Aegilops* pop 2” batch, and the “Wheat pop” batch, representing *A. cylindrica*, *A. tauschi* and *T. aestivum* derived isolates, respectively. These batches of isolates were inoculated individually on to different wheat and wild grasses accessions (S4 Table) along with a negative “mock” control and a “positive” control represented by the reference *Z. tritici* isolate IPO323. The disease symptoms were recorded at 21 and 28 days post-inoculation (dpi). Out of the 31 different wheat and wild-grass relatives tested, only *A. cylindrica* could be infected by the isolates present in the “*Aegilops* pop 1” batch. No symptoms were observed for any *Triticum* species inoculated with this batch (S11 Fig and S5 Table). The isolates contained in the “*Aegilops* pop 2” batch did not produce symptoms in any of the *Triticum* species used in the experiment but were virulent on *Aegilops cylindrica* and *Aegilops tauschii* plants (S11 Fig and S5 Table). Isolates belonging to the “Wheat pop” batch were able to infect *Triticum aestivum* and other closely-related *Triticum* species (e.g. *T. spelta* and *T. compactum*) in addition to *Aegilops tauschii* (S11 Fig and S5 Table).

Considering the clear virulence dichotomy on *A. cylindrica* observed between isolates from the *Aegilops* population 1 and the wheat population, we performed a second qualitative infection assay focusing just on these two populations. In this experiment, we aimed to confirm the extent of host specificity among the collection of sequenced isolates from the *Aegilops* population 1 and the wheat population by selecting two to three isolates per herbarium sample and inoculating them individually on *A. cylindrica* and *T. aestivum* plants. Consistent with the previous inoculation assay, *Aegilops*-infecting *Z. tritici* isolates could only infect *A. cylindrica* plants, while the wheat-infecting isolates could only cause symptoms on *T. aestivum* plants (Fig 3A and S6 Table). Taken together, these results support our hypothesis that host specificity is shaping the population structure of these sympatric yet divergent *Z. tritici* populations and reinforce the assumption of different host-specific *Z. tritici* lineages. In addition, we could confirm the virulence of individual *Z. tritici* isolates on *A. cylindrica* plants and thereby establish an experimental system for the *Aegilops*-infecting *Z. tritici* pathogen.

**Fig 3.**
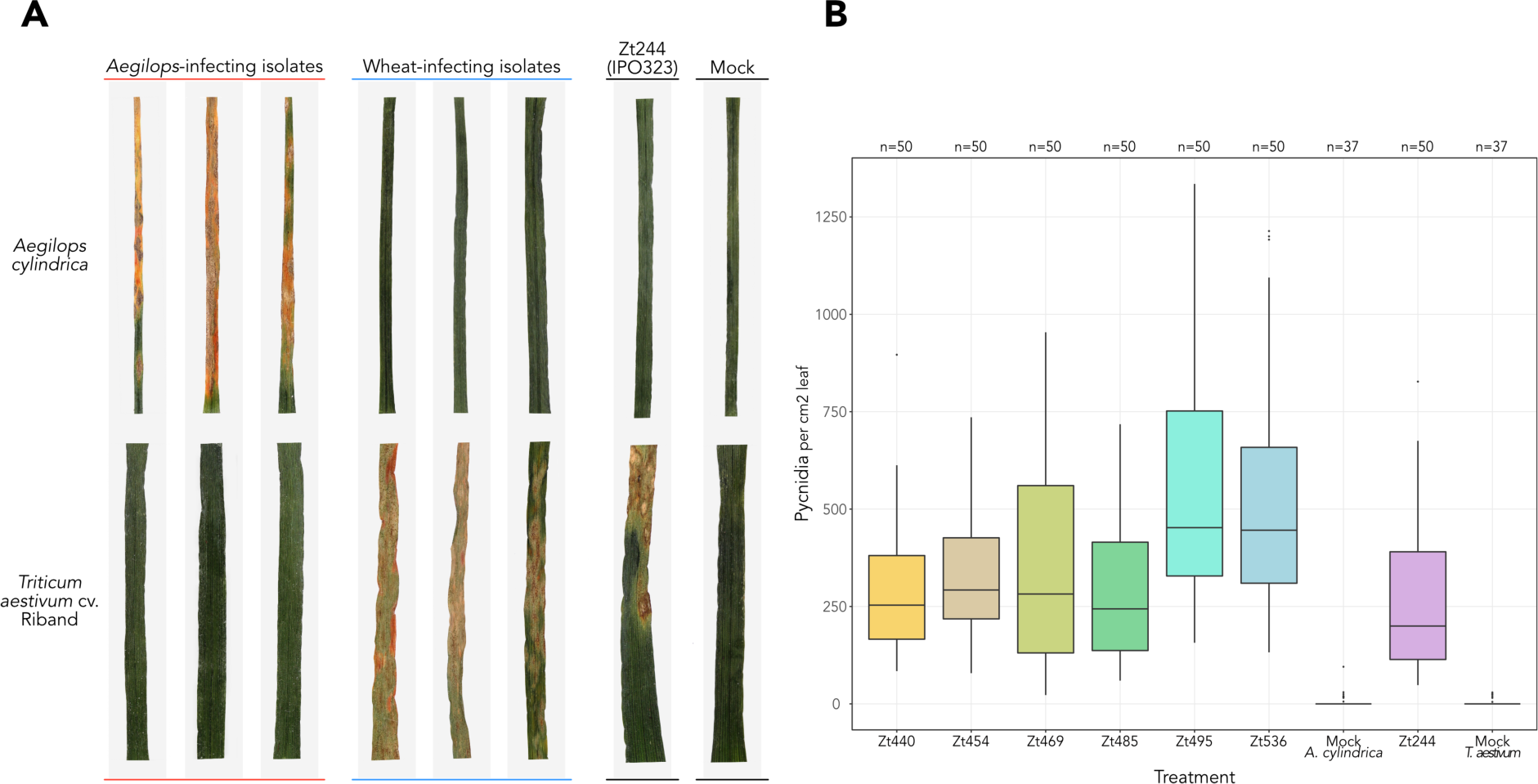
Host specificity and quantitative virulence among *Z. tritici* isolates. **(A)** Summary of qualitative virulence assays performed with *Z. tritici* isolates in the greenhouse. *Aegilops cylindrica* (top panels) and wheat (*Triticum aestivum* cv. Riband; bottom panels) were inoculated with adjusted blastospore suspensions of each individual *Z. tritici* isolate. Each grey column represents leaves inoculated with a different isolate. Mock plants were treated with 0.1% Tween® 20 in sterile water. Pictures were taken at 21 dpi (days-post inoculation). Complete description of qualitative virulence results can be found in S6 Table. **(B)** Quantitative analyses of *Z. tritici* virulence in distinct hosts. Leaves of *A. cylindrica* were inoculated with a subset of *Aegilops*-infecting *Z. tritici* isolates from *Aegilops* population 1. As a comparison to a wheat-infecting *Z. tritici* isolate, *T. aestivum* cv. Riband plants were inoculated with the *Z. tritici* IPO323 isolate (strain Zt244). Leaves were scanned at 21 dpi and pycnidia density (per cm^2^ leaf) was used as a readout of virulence. Number of leaves per treatment (n) are indicated. Kruskal-Wallis test indicates statistical difference in pycnidia densities among all treatments tested (chi-squared = 218.08, df = 8, P-value < 2.2e-16).

In order to better characterize the *Z. tritici* infection development in the *A. cylindrica*-*Z. tritici* pathosystem, we performed a disease progression analysis and conducted a quantitative measure of disease symptoms. In these assays, we used a subset of *Aegilops*-infecting *Z. tritici* isolates from *Aegilops* population 1 along with the *Z. tritici* IPO323 isolate (strain Zt244) and mock treatments as positive and negative control, respectively. All isolates/treatments were inoculated individually at similar inoculum concentrations on the correspondent hosts: *A. cylindrica*, for the *Aegilops*-infecting *Z. tritici* isolates and *T. aestivum* cv. Riband, for the *Z. tritici* IPO323 isolate. Disease progression analyses revealed that leaves inoculated with the *Aegilops*-infecting *Z. tritici* isolates showed, on average, the first signs of necrosis and pycnidia at 9-11 dpi and 11-13 dpi, respectively (S12 Fig). This onset of visible symptoms was earlier than those observed for the wheat-infecting *Z. tritici* isolate Zt244 (IPO323), in which visible symptoms became apparent between 15 dpi to 19 dpi on *T. aestivum* plants (S12 Fig). Similar results for other wheat-infecting isolates were observed previously, with necrosis and pycnidia becoming visible 12-19 dpi and 17-23 dpi, respectively [56].

Next, we performed a quantitative measure of disease symptoms. To this end, we inoculated *A. cylindrica* and wheat plants with the same isolates used for the disease progression analyses and collected inoculated leaves at 21 days post-inoculation (dpi) to evaluate infections using pycnidia density as a measure of virulence. We observed statistically different levels of pycnidia among all treatments tested (Kruskal-Wallis test, chi-squared = 218.08, df = 8, P-value < 2.2e-16; Fig 3B). The isolates Zt495 and Zt536 showed higher pycnidia densities compared to the other isolates, although no statistical difference was observed between these two isolates (Wilcoxon Rank Sum test, P-value = 0.65; Fig 3B). Altogether, the differences in the temporal disease development and in the levels of virulence observed among the *Aegilops*- and wheat-infecting isolates suggest that the infection development of *Z. tritici* can vary in pace in different grass hosts.

### Infection of *Z. tritici* in *A. cylindrica* is characterized by four developmental stages

Next, we addressed if the infection development of *Z. tritici* isolates on *Aegilops cylindrica* plants was similar to the one previously observed for *Z. tritici* on wheat [56]. To this end, we morphologically characterized Zt469 host colonization using detailed confocal scanning laser microscopy (CSLM) analyses in inoculated *A. cylindrica* leaves collected between 7 and 21 dpi (see Methods). The isolate Zt469 was randomly selected to represent the *Aegilops* population 1 and it was included in all population genomics and virulence assays datasets analyzed in this study. Leaf samples were characterized based on the previously described infection development stages of *Z. tritici* on wheat [56] in a single-blinded procedure (see Methods). Briefly, we characterized infection events belonging to the following four developmental stages: stage “A”, or infection establishment stage, in which fungal hyphae penetrate leaf tissue via stomata; stage “B”, characterized by biotrophic, symptomless colonization of leaf mesophyll by fungal hyphae; stage “C”, which comprises the transition from biotrophic to necrotrophic growth when first disease symptoms are apparent; and the stage “D”, characterized by necrotrophic colonization and asexual reproduction. In this stage, mesophyll tissue is heavily colonized by hyphae and substomatal cavities are occupied by mature pycnidia harboring asexual pycnidiospores [56]. Based on these characteristics, we identified leaf samples which represented these four infection stages, namely 7dpi (stage “A”), 10dpi (stage “B”), 15dpi (stage “C”) and 21dpi (stage “D”; S13 Fig). These results indicate a conserved infection program of *Z. tritici* on *Aegilops* and *Triticum* hosts.

### Host-diverging *Z. tritici* populations exhibit distinct selective sweep footprints

Based on the findings that the analyzed sympatric *Z. tritici* populations show distinct clusters based on genomic data and moreover exhibit host specificity in inoculation experiments, we set out to investigate which traits may be responsible for the different host specificities. We hypothesized that genetic traits responsible for host specificity would exhibit signature of recent positive selection. We therefore performed genome scans to detect signatures of selective sweeps in each individual population. For these analyses, we focused on the *Aegilops* population 1 and the wheat-infecting population. Our analyses were restricted to SNPs for which we could assign the ancestral states from a comparison of genome-wide *Z. tritici* SNP data to the corresponding sites in *Z. ardabiliae* and *Z. pseudotritici*. Ancestral SNP alleles were designated if the same SNP alleles was fixed in both sister species. Using this approach, our selective sweep scans were based on 149,716 SNPs with known ancestral states dispersed over the thirteen core chromosomes of *Z. tritici*.

Signals of selection can be erroneously detected in genome-wide scans. Demographic events as population bottlenecks or migration events, in particular, can introduce confounding factors to the analyses and result in false positives [23,24,57]. To decrease the number of false positives, we used two methods and stringent outlier thresholds to detect selective sweeps: the Composite Likelihood Ratio (CLR) method, implemented in SweeD [58]; and the μ statistic, implemented in the RAiSD software [59], which detects different signatures of selective sweeps across chromosomes. These two methods slightly differ in terms of detection of hard selective sweeps (i.e. selection and spread of a single advantageous allele in a population; [60,61]) while being robust to demographic scenarios [23,24,57,59]. After clustering sweep regions based on high 99.5% outlier thresholds and linkage disequilibrium distances (see Methods), several and widespread selective sweep regions were detected across all chromosomes among the host-diverging populations (Fig 4 and S7-S8 Tables). Selective sweep regions were on average 25066 bp and 8100 bp in length in the *Aegilops* and wheat-infecting *Z. tritici* populations, respectively (S8 Table). This difference may reflect the reduced extent of linkage disequilibrium in the wheat-infecting population providing a higher resolution of sweep regions. We also observed that some selective sweep regions were commonly detected by the two methods within each population. In the *Aegilops* population 1, we found two regions (in chromosomes 4 and 7) while five regions (in chromosomes 1, 5 and 9) were in common between the two methods in the wheat population (Fig 4 and S7 Table). Interestingly, different regions have been under selection in the wheat-infecting and *Aegilops*-infecting populations with only one region in chromosome 5 overlapping between the two *Z. tritici* populations (S7 Table).

**Fig 4.**
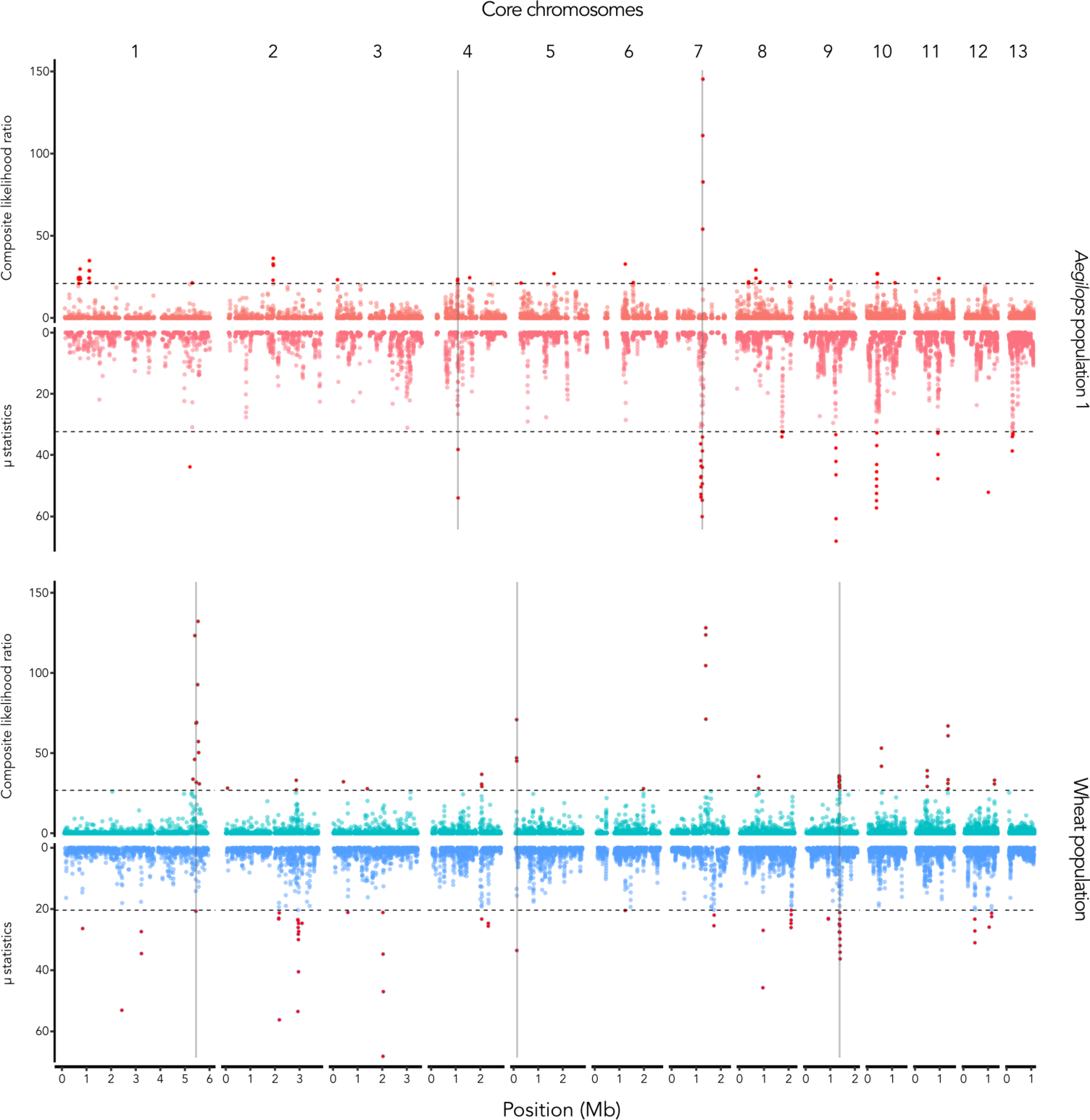
Diverse and widespread selective sweep signatures in host-diverging *Z. tritici* populations. Genome scans for selective sweeps were performed across the thirteen core chromosomes of *Z. tritici*. Two methods were used for each Iranian host-diverging *Z. tritici* population. Top panels show the CLR and μ statistics scores for the *Aegilops* population 1 while the bottom panels show the same statistics for the wheat population. Horizontal dashed lines represent the 99.5th quantile threshold in each method and individual population. Above the thresholds, outlier regions potentially under selective sweep are highlighted in red. Selective sweep regions commonly detected by the two methods within each population are represented by vertical grey bars.

### Selective sweep regions harbor candidate genes for host adaptation

To investigate genes that are putatively under recent selection and potentially involved in host adaptation, we characterized the genetic composition of selective sweep regions in each population individually. We used the gene and Transposable Element (TE) models and functional annotations in the reference *Z. tritici* isolate IPO323 published previously [62–64] to identify the features present in the regions under selection. In total, selective sweep regions harbored 301 genes and 184 genes in the *Aegilops* and wheat-infecting *Z. tritici* populations, respectively (S9 Table). Regarding TEs, selective sweep regions contained a total of 43 TEs for the *Aegilops* population 1 and 46 TEs in the wheat population (S9 Table). We also performed Gene Ontology (GO) analyses and tested the enrichment of GO terms and PFAM domains in the selective sweep regions compared to whole genome. GO terms could be assigned to all *Z. tritici* genes analyzed in both populations while 69% (208/301) and 66% (121/184) of the genes under selective sweep had annotated PFAM domains in the *Aegilops* population 1 and wheat population, respectively. For the *Aegilops* population 1, we observed that GO terms for biological processes related to RNA splicing and DNA replication and transcription were overrepresented among the selective sweep regions (Fischer’s exact test, P-values = 0.0089 - 0.045; S10 Table). In the selective sweep regions of the wheat population, we found that GO terms related to biosynthetic processes and transport were significantly enriched (Fischer’s exact test, P-values = 0.029 - 0.049; S10 Table). PFAM domain analyses furthermore revealed that in the wheat population, selective sweep regions were enriched in genes encoding cytochrome P450-like and polyketide synthase-like proteins, chitin-binding lysin motif (LysM), among others (χ2 test, P-value < 0.05; S11 Table). In the *Aegilops* population 1, we observed a significant enrichment of genes potentially involved in pathogenicity and encoding glycosyl hydrolases, peptidases, antibiotic biosynthesis monooxygenase along with other diverse domains (χ2 test, P-value < 0.05; S11 Table).

In host-pathogen interactions, different pathogen virulence factors, including genes encoding effectors and carbohydrate-degrading enzymes (CAZymes), are important components of the molecular arms race with host plants [13,65]. Focusing on these gene categories, we observed that five candidate effector genes were present in the selective sweep regions of the *Aegilops* population 1 while two were identified in the wheat population (S9 Table). The selective sweep regions in *Aegilops* population 1 comprised thirteen CAZymes-encoding genes while in the wheat population nine CAZymes-encoding genes were identified (S9 Table). None of the effector or CAZyme-encoding genes present in the selective sweep regions were in common between the two host-diverging *Z. tritici* populations. Interestingly, among the different selective sweep regions in the *Aegilops* population 1, one region stood out. We found that from all the five candidate effector genes detected in the selective sweep regions of the *Aegilops* population 1, three candidate effectors (Zt09_chr_7_00299, Zt09_chr_7_00305 and Zt09_chr_7_00308) were localized in proximity at chromosome 7, spanning a unique selective sweep region identified from position 1,000,062 to 1,032,926bp using the CLR test (S7 and S9 Table). Moreover, adjacent to this selective sweep region, RAiSD also identified a region potentially under selection from position 980,772 to 1,013,262bp (S7 and S9 Table). Part of this genomic region (from 1,000,062 bp to 1,013,262 bp in chromosome 7) was the one observed to be in common between the two selective sweep metrics used for the *Aegilops*-infecting population containing one of the candidate effector genes (Zt09_chr_7_00299) (Fig 4, S7 and S9 Table).

Further considering the detection overlap and the importance of effectors and CAZymes in host-pathogen interactions, we conducted a more detailed characterization of the selective sweep region identified on chromosome 7 in the *Aegilops* population 1. We used the genomic coordinates detected by the CLR test as they comprised the three candidate effector genes within the same region. In this local analysis, we also included part of the selective sweep region identified by RAiSD (which encompasses the localization of the candidate effector gene Zt09_chr_7_00299) and surrounding genomic locations by adding 15 kb to the up and downstream flanks of the selective sweep region defined by the CLR test. Moreover, to compare the pattern of genetic diversity between the two host-diverging *Z. tritici* populations, we analyzed the region using SNPs from the wheat population and *Aegilops* population 1 individually. Intriguingly, we observed peaks of high absolute (Dxy) and relative (Fst) divergences between the *Aegilops*- and wheat-infecting populations across the region under selection, particularly in the loci encoding the three candidate effector genes (Fig 5A, B). We moreover observed a lower nucleotide diversity (*π*) for the *Z. tritici* isolates infecting *Aegilops* spp. throughout the whole selective sweep region as well as in the surrounding flanking regions (Fig 5A). A decrease in nucleotide diversity was also observed in the wheat-infecting population for the regions encoding the three candidate effector genes (Fig 5A). We also identified negative Tajima’s D values in the *Aegilops* population 1, specifically in the region comprising the CAZyme gene (Zt09_chr7_00296) and two candidate effector genes (Zt09_chr7_00299 and Zt09_chr7_00305), while the isolates infecting wheat showed a positive Tajima’s D value in the detected selective sweep locus (Fig 5A, B). We found a high linkage disequilibrium between pairs of SNPs across the region detected in the selective sweep scan for the *Aegilops*-infecting *Z. tritici* isolates, particularly in the region spanning the genes Zt09_chr7_00296 and Zt09_chr7_00308 (Fig 5C). In contrast, the wheat-infecting population showed a low linkage disequilibrium throughout the whole selective sweep region and surrounding flanking regions (Fig 5C). To complement these results, we performed the McDonald & Kreitman (MK) test for each gene within the selective sweep region using the polymorphisms of each host-diverging population individually, and *Z. ardabiliae* gene orthologs as outgroups. No significant differences in the ratio for synonymous versus non-synonymous sites within and between species was detected in any gene analyzed in the *Aegilops* population 1. In the wheat-infecting population, evidence for non-neutral evolution was however found for one gene, namely Zt09_chr_7_00300 (P-value < 0.001).

**Fig 5.**
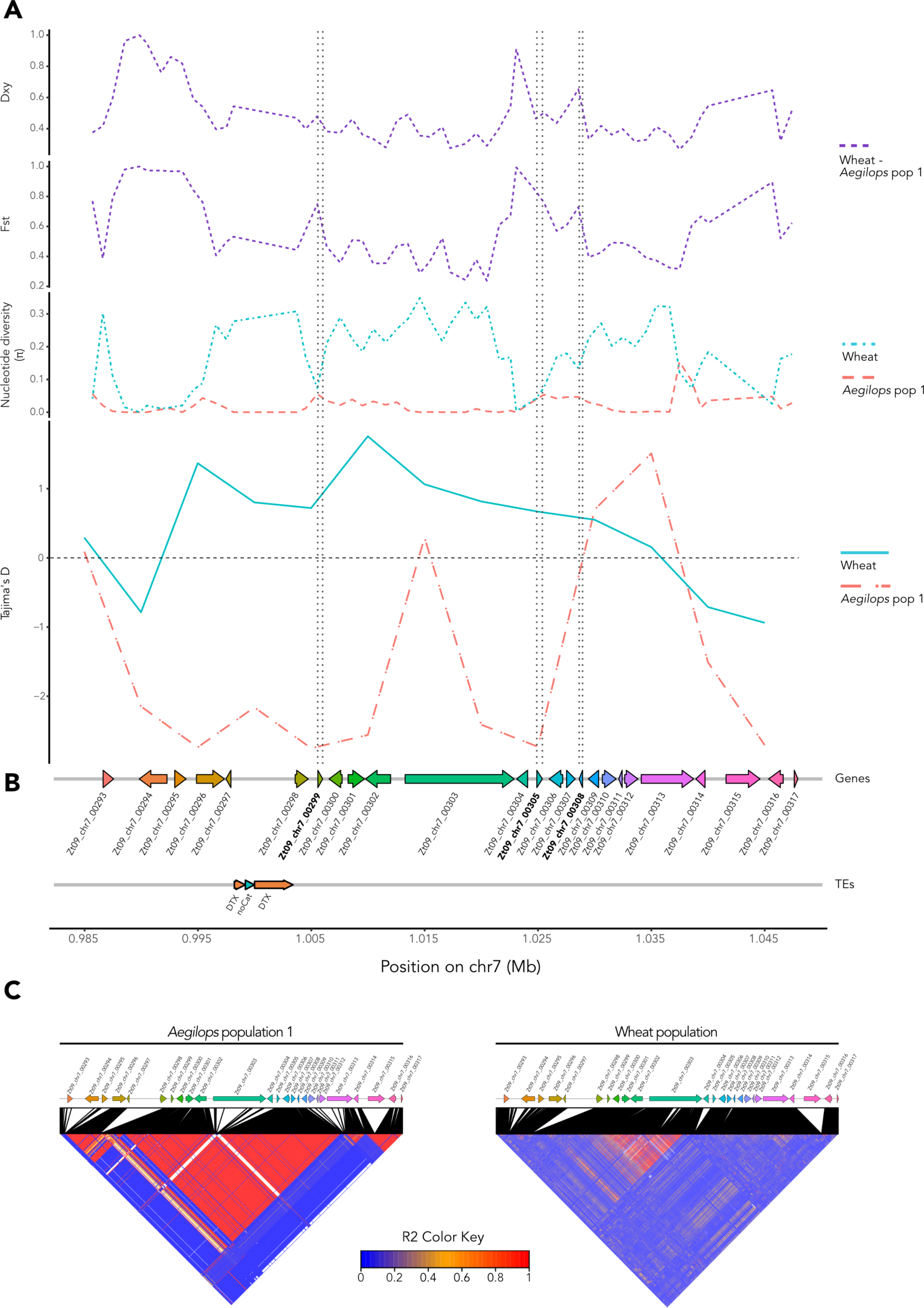
*Aegilops* population 1 shows selective sweep signatures at chromosome 7. Genomic summary statistics were calculated across the selective sweep region with 15kb up- and downstream flanks for the *Aegilops* population 1 and wheat population individually. **(A)** Absolute (Dxy) and relative (Fst) divergences between the two host-diverging *Z. tritici* populations (top panels) and nucleotide diversity (*π*) for each population (middle panel) were calculated in 1kb windows. Tajima’s D (lower panel) was calculated in 5kb windows. Vertical dashed lines delimit the genomic locations of the three candidate effector genes. **(B)** Genes and TE models annotated across the selective sweep and surrounding genomic regions (DTX = Class II transposons, TIR; noCat = no category). Candidate effector genes are in bold. **(C)** Linkage disequilibrium heatmap based on coefficient of correlation (r2) between SNPs in the *Aegilops* population 1 (left) and wheat population (right). Correspondence between SNPs location in the genomics regions (e.g. genes) and on the heatmap are depicted by black segments.

Taken together, these results show clear patterns of selection in the selective sweep region of chromosome 7, even though the MK tests did not detect significant signals of positive selection. The negative Tajima’s D as well as the high linkage disequilibrium values observed for the *Aegilops* population 1, in particular, support the signatures of a recent selective sweep at this locus [23,24,66]. More importantly, the high peaks of absolute (Dxy) and relative (Fst) divergence detected between the two host-diverging population on candidate effectors further pinpoint candidate genes that may contribute to host adaptation of these distinct populations.

### Candidate effector genes in selective sweep show temporal-specific expression during *A. cylindrica* infection

Next, to gain further insights into the potential relevance of the three candidate effector genes in the selective sweep region of chromosome 7 for pathogenicity in *A. cylindrica*, we analyzed their expression patterns in Zt469 during *in vitro* growth and *A. cylindrica* infection. As a comparison, we also analyzed the expression patterns of the homologous candidate effector genes in the *Z. tritici* isolate IPO323 during *in vitro* growth and during wheat infection. To this end, we used previously generated RNA-seq datasets of these *Z. tritici* isolates during growth on YMS medium and during infection at four different stages in *A. cylindrica* (for Zt469; Fagundes et al. 2024) and in wheat (for IPO323; [56]). In the *Z. tritici* IPO323 genome, analyses of the normalized abundance of transcripts in FPKM (Fragments per Kilobase of Transcript per Million fragments mapped) revealed that the three candidate effectors had higher levels of expression during wheat infection when compared to *in vitro* growth (S14 Fig). The candidate effector gene Zt09_chr_7_00299 had a constant increase of expression with the progression of infection from 4 days-post inoculation (dpi) to 20 dpi as previously reported (S14 Fig; [56]). The other two candidate effector genes Zt09_chr_7_00305 and Zt09_chr_7_00308 were not expressed during *in vitro* growth on YMS medium, while *in planta* they were both expressed at much lower levels than Zt09_chr_7_00299 (S14 Fig; [56]). Interestingly, these two candidate effector genes did not have a detectable expression at the infection stage “B” on wheat (S14 Fig).

In the *Aegilops*-infecting *Z. tritici* isolate Zt469 however, we observed distinct expression patterns for the three homologues candidate effector genes (S14 Fig). For the candidate effector gene Zt469_000007F_arrow_0295 (homolog: Zt09_chr_7_00299), we also observed higher levels of expression *in planta* than *in vitro*, however with a peak of expression at stage “B” and a constant reduction of expression at later stages (“C” and “D”). The candidate effector gene Zt469_000007F_arrow_0301 (homolog: Zt09_chr_7_00305) only showed expression during *in vitro* growth and at stage “D” while the candidate effector gene Zt469_000007F_arrow_0302 (homolog: Zt09_chr_7_00308) only showed expression during *in vitro* growth (S14 Fig). Altogether, these results suggest that the candidate effectors genes found in the selective sweep region of chromosome 7 have a temporal-specific expression regulation during *in planta* infection in both *A. cylindrica*- and wheat-*Z. tritici* pathosystems.

### Deletion of one candidate effector gene in selective sweep impacts virulence of *Aegilops*-infecting *Z. tritici*

Considering the high expression levels observed for the candidate effector gene Zt09_chr_7_00299 during *in planta* infection and its detection in both selective sweep scans in *Aegilops* population 1 (S14 Fig, S7 and S9 Tables), we hypothesize that this gene plays a relevant role for *Z. tritici* pathogenicity on *A. cylindrica* plants. To test this hypothesis, we generated knock-out mutants for this candidate effector gene using the *Aegilops*-infecting *Z. tritici* Zt495 from *Aegilops* population 1. We chose the *Z. tritici* isolate Zt495 as a background for such deletion given its high virulence in *A. cylindrica* plants (Fig 3B). Deletion mutants were generated using an *Agrobacterium tumefaciens*-mediated transformation (ATMT) approach ([67]; see Methods) and the correct integration of the selection marker cassette by homologous recombination was confirmed by PCRs and Southern blot analyses in three independent mutant strains (strains #68, #86 and #98).

Then, in order to test the impact of the candidate effector gene deletion on virulence, we set out a quantitative virulence assay under greenhouse conditions using *A. cylindrica* as a host. We inoculated *A. cylindrica* plants with adjusted blastospore suspensions of the Zt495 wild-type and the three deletion mutant strains. Mock treatments were also inoculated on *A. cylindrica* plants and served as negative controls. Analyses of pycnidia density in leaves collected at 21 dpi revealed a statistical difference in pycnidia levels between each of the deletion mutant strains and the Zt495 wild-type isolate, with lower levels detected for the mutant strains (pairwise Wilcoxon Rank Sum test and Holm’s correction, P-value < 0.05; Fig 6). These results indicate that the candidate effector gene Zt09_chr_7_00299 plays an important role for virulence of *Z. tritici* on *A. cylindrica* plants and hence the genomic region encoding the gene has recently undergone a selective sweep.

**Fig 6.**
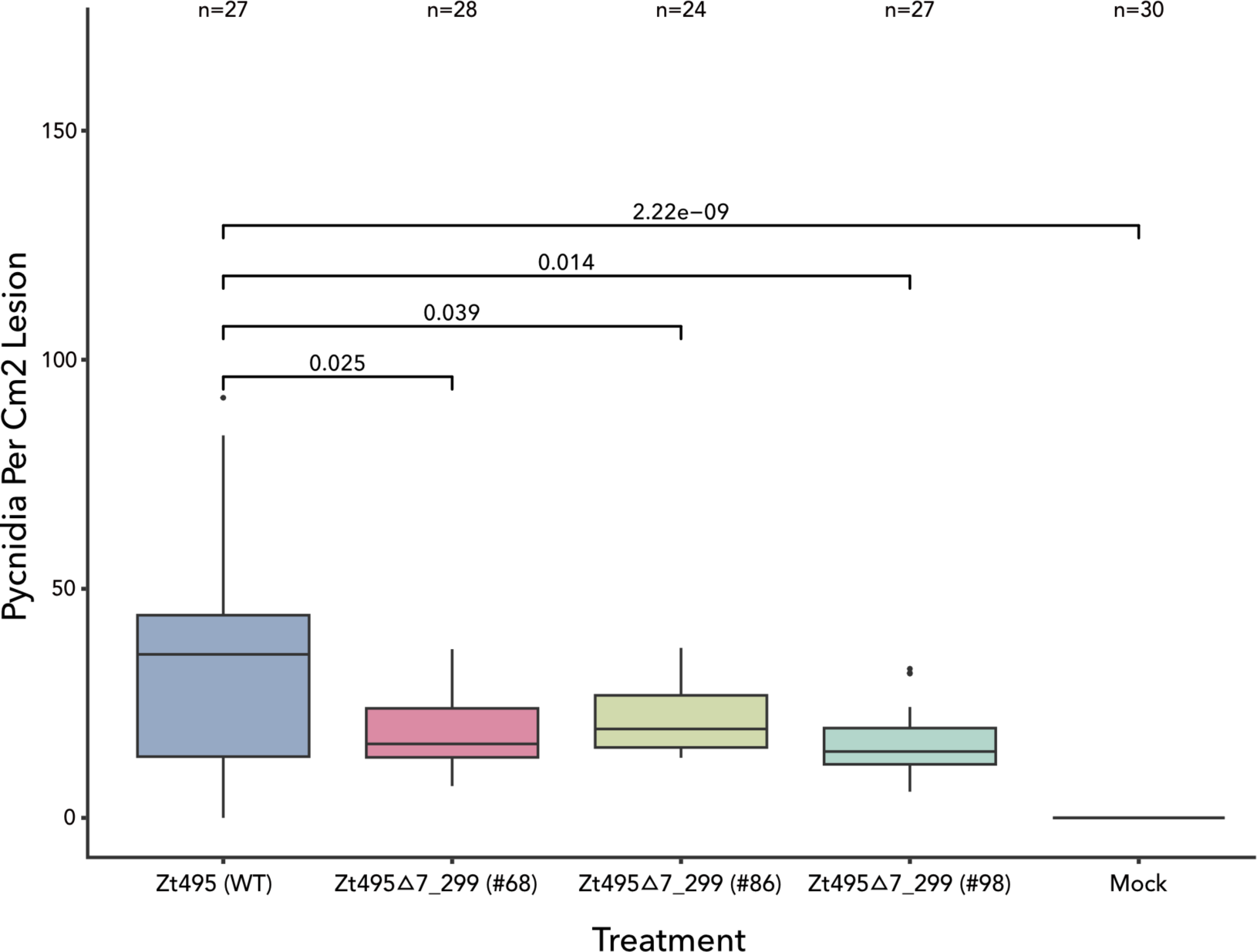
Knock-out *Z. tritici* mutants of candidate effector gene in selective sweep show reduced virulence on *A. cylindrica* plants. Leaves of *A. cylindrica* plants were inoculated with adjusted blastospore suspensions of the wild-type Zt495 isolate and of the three deletion mutant strains (#68, #86 and #98) for the candidate effector gene Zt09_chr_7_00299 (“Δ7_299”). Mock treatment was used as a negative control. Leaves were collected at 21 dpi and pycnidia density (per cm^2^ lesion) was measured as a readout of virulence. Numbers over brackets represent p-values of pairwise Wilcoxon rank-sum tests and Holm’s correction for multiple testing. Number of leaves per treatment (n) are indicated.

## Discussion

Host adaptation is a necessary step in the evolution and divergence of new pathogen species [3,4,68]. Although many pathogens exhibit strong host specificity, only few studies have identified traits that determine this specificity. In this study, we present comprehensive analyses on unique host-diverging *Z. tritici* populations and further demonstrate the impact of host adaptation on fungal evolution and population divergence. Using population genomics analyses and comparative infection assays, we show that host specificity is a strong determinant of *Z. tritici* population structure and identify candidate genes likely involved in host adaptation of this important fungal pathogen.

Previous studies have demonstrated that *Z. tritici* populations are highly diverse within wheat fields and show a strong continental population structure consistent with the geographic origin of the isolates [28,33,34,69]. Host cultivar specificity has also been shown to be an important factor in the interaction between wheat and *Z. tritici* isolates [70–76]. Mechanisms of *Z. tritici* host specialization have also been studied in durum wheat (*T. durum*) [73,75,77–79] and in the noncompatible grass host *Brachypodium distachyon* [80]. However, while these studies have shown important factors underlying the interaction between *Z. tritici* and specific hosts, none have unraveled population structure and genome evolution on distinct host species or host genotypes. Here we show that sympatric *Z. tritici* isolates, derived from the center of origin of these pathogens, cluster in three distinct populations which corresponds to host species. We find few signatures of admixture between *Z. tritici* populations on wild and domesticated wheat species. Corroborating the genomics analyses, we show, by barcode-amplicon sequencing of herbarium specimens, that different host species are indeed the determinants of the population structure in the Iranian *Z. tritici* populations and further confirm their host specificity through infection assays.

We observed distinct patterns of linkage disequilibrium (LD) and differences in the overall genetic diversity in our host-diverging *Z. tritici* populations. These results suggest that besides being genetically distinct, these populations also differ in the extent of sexual versus asexual propagation. In line with this, we also observed that these populations differ in the frequencies of mating type idiomorphs. During the process of adaptation to a new host habitat, fungi can undergo frequent asexual propagation with rare events of sexual reproduction [3,81]. The lack of sexual recombination can still enhance the accumulation of genomic variants by genetic drift, which in turn can act as barriers of sexual recombination and gene flow with genetically diverged individuals from other host habitats and therefore continuously increase the divergence between these populations [3,4,19]. Thus, while our observations may be a reflection of the sampling we used, they also suggest that specialization on different host can impact the frequency of sexual reproduction in *Z. tritici* and thereby may contribute to the divergence we observe between the different host-infecting *Z. tritici* populations.

Previous comparisons between pathogens occurring on domesticated and wild host species have found contrasting patterns. In the plant pathogenic fungi *Rhynchosporium secalis* [82] and the apple scab fungal pathogen *Venturia inaequalis* [83] specialization on a crop host has entailed loss of variation, while genetic diversity in *Z. tritici* infecting wheat has increased. Here, we also find that the *Aegilops*-infecting *Z. tritici* populations exhibit a lower genetic diversity compared to the domesticated wheat-infecting population. We hypothesized that this is due to two factors: (i) large effective population sizes in the domesticated pathosystem and (ii) a stronger effect of fluctuating population sizes in the wild pathosystem involving recurrent bottlenecks. On one hand, the dense and uniform host populations found in agricultural ecosystems allow pathogen populations to rapidly propagate, and in the case of sexually reproductive species, increase the genetic diversity of the local population by recurring cycles of reproduction throughout the season [68]. This increase in population size directly impacts the maintenance of genetic diversity given that populations with a larger population size are less prone to genetic drift [84]. In the case of *Z. tritici* on wheat, several population genomics studies have shown that the mixed reproductive system found in the *Septoria tritici* blotch (STB) epidemics (characterized by both asexual and sexual reproduction) allows this fungus to maintain high levels of both gene and genotype diversity in field populations around the world [33,34,85–90]. In fact, a recent study estimated that wheat fields showing typical STB infections can carry up to 14.0 million *Z. tritici* genotypes per hectare, indicating a large evolutionary potential of the local *Z. tritici* populations and maintenance of high genomic diversity [91]. On the other hand, host populations in wild ecosystems generally consist of genetically diverse individuals with a heterogeneous distribution in space and time [10]. This sparse distribution of host populations leads to fluctuations in the local pathogen population sizes and contribute to the metapopulation dynamics of these pathogens on wild plants [92,93]. The metapopulation dynamics of local pathogens in wild ecosystems is characterized by recurrent cycles of population extinction and re-colonization, leading to frequent population bottleneck events and a severe loss of genetic variation at local scales [10,94,95]. Considering the annual and variable distribution of *Aegilops* species in different ecosystems and climates throughout Iran [96], the metapopulation dynamics can be a strong factor contributing to the low genetic diversity found in the *Aegilops*-infecting *Z. tritici* populations.

Adaptation to distinct hosts has impacted not only the population structure but has also leave footprints of recent positive selection in genomes. We performed genome scans using the CLR and μ statistics methods and stringent thresholds to detect selective sweeps in each host-diverging *Z. tritici* population. Regions under recent positive selection were widespread and diverse among the populations analyzed with only one region overlapping between the *Aegilops* and the wheat-infecting *Z. tritici* populations. The observed differences in the number and location of selective sweeps indicate that these host-diverging populations have experienced selection in different regions of their genomes. A previous study analyzing selective sweeps across the *Z. tritici* genome has also found differences in the number and position of selected regions among four allopatric wheat field populations [28]. The authors argued that differences in fungicide usage, annual mean temperatures and deployed host cultivars in the fields are likely factors contributing to such heterogeneous selection observed between the analyzed populations [28,69,97–99]. Regarding wild ecosystems, other studies have pointed out that divergent selection can act upon fungal populations isolated across distinct ecological niches and leave signatures of positive selection at different loci [25–27]. Thus, we hypothesize that the distinct selective sweep regions detected between the *Aegilops*- and wheat-infecting *Z. tritici* populations can be explained by the divergent evolutionary history of these populations and the concomitant local adaptation of these isolates to their host environment, as exemplified by our demography and population structure analyses.

In the wheat population, we observed that selective sweep regions were enriched in genes encoding polyketide synthase-like proteins and chitin-binding lysin motifs (LysM; S11 Table). Polyketide synthase genes have been shown to be upregulated in *Z. tritici* during wheat infection and suggested to play a role during disease progression by producing yet unknown secondary metabolites targeting the host (e.g. toxins) or members of the host microbiome [100,101]. Many fungal pathogens have evolved effector proteins containing LysM motifs that can either protect their cell walls against plant chitinases or prevent the recognition and elicitation of chitin-triggered host immunity [102–109]. In *Z. tritici*, the LysM effectors Mg1LysM and Mg3LysM have been identified and functionally characterized, and more recently the crystal structure of the effector Mg1LysM was elucidated [104,110]. These analyses have revealed that Mg1LysM and Mg3LysM have the ability to protect fungal hyphae against host chitinase hydrolysis and contribute to fungal virulence.

In the *Aegilops* population 1, we observed that selective sweep regions were enriched with genes encoding peptidases, diverse glycosyl hydrolases (GHs), and antibiotic biosynthesis monooxygenases (Abm; S11 Table). In fungal pathogens, secreted peptidases can act as virulence factors and contribute to pathogenesis, as already demonstrated in the fungal pathogens *Botrytis cinerea* [111], *Fusarium culmorum* [112], *Glomerella cingulata* [113] and *Sclerotinia sclerotiorum* [114]. Glycosyl hydrolases (GHs), which represent the largest class of carbohydrate-active enzymes (CAZymes) and often act as plant cell wall degrading enzymes (PCWDEs), have also been shown to be an important group of secreted proteins contributing to fungal virulence and endophytic lifestyle [115–120]. However, analyses of the reference *Z. tritici* IPO323 genome have revealed a small complement of PCWDEs compared to other plant pathogenic fungi [121,122] and a low expression of these genes during the biotrophic phase of wheat infection [17], suggesting a biphasic mechanism of nutrient-stealth pathogenesis in the host apoplast as proposed by Goodwin and collaborators [121]. The presence of one gene encoding antibiotic biosynthesis monooxygenases (Abm; PFAM PF03992) within the selective sweep region on chromosome 7 of the *Aegilops*-infecting population also points to new candidate genes involved in virulence (S9 Table). A study by Patkar and collaborators (2015) showed that the rice blast fungus *Magnaporthe oryzae* uses the antibiotic biosynthesis monooxygenase (Abm) to convert intrinsically produced as well as host-derived JA into 12-hydroxyjasmonic acid (12OH-JA) during infection, which in turn suppresses JA-mediated signaling defense responses in the rice host and thus facilitates host colonization [123,124]. This plethora of genes and gene functions detected between the two host-diverging *Z. tritici* populations suggests that adaptation to distinct hosts drives selection on a wide range of traits targeting several host defense mechanisms.

Considering the importance of effectors for pathogenesis and host adaptation [13,65,125,126], we particularly focused on the distribution of genes encoding candidate effector proteins throughout the selective sweep regions. Interestingly, we found a specific selective sweep region on chromosome 7 in the *Aegilops*-infecting population harboring three candidate effector genes in close proximity (Fig 5). Adjacent to these genes, we also found a CAZyme-encoding gene (Zt09_chr_7_00296) and one of the genes with the antibiotic biosynthesis monooxygenase (Abm) domain (Zt09_chr_7_00297). Our more detailed analyses on the genomic landscape in this selective sweep region revealed remarkable footprints of recent positive selection and high divergence between the two host-diverging *Z. tritici* populations.

A previous study analyzing isolate- and temporal-specific infection programs of *Z. tritici* in wheat has shown the expression of particular candidate effector genes and pointed out their potential contribution for virulence in different infection stages [56]. In this study, Haueisen and colleagues (2019) demonstrated that in the *Z. tritici* isolate IPO323 (strain Zt09), the candidate effector gene Zt09_chr_7_00299 was upregulated during wheat infection, and this expression had a constant increase from 4 days-post inoculation (dpi) to 20 dpi [56]. In our study, we showed that the homologous candidate effector gene in *Aegilops*-infecting *Z. tritici* Zt469 (Zt469_000007F_arrow_0295) also shows high expression levels during *in planta* infection however with a different temporal pattern (S14 Fig). Using knock-out mutant strains, we further confirm a pathogenicity role of this candidate effector gene in *Aegilops*-infecting *Z. tritici.* Hence, these results indicate that the detected selective sweep region in chromosome 7 harbors genes involved in *Z. tritici* virulence and potentially host adaptation. Complementation strains and “allele-swap” mutants for the candidate effector gene Zt09_chr_7_00299 between wheat- and *Aegilops*-infecting *Z. tritici* strains are important next steps to confirm and elucidate the role of this gene in pathogenesis and host specificity.

A previous phylogenetic study in *Zymoseptoria* species has pointed out the presence of a host-diverging phylogenetic cluster within the *Z. tritici* clade collected from *Aegilops tauschii* plants in Iran [127]. In agreement with Quaedvlieg and collaborators (2011), we here also find evidence for the genetic differentiation of different *Z. tritici* populations on different host species; a signature that may reflect incipient speciation.

*Zymoseptoria tritici* is considered a specialized pathogen of wheat although reports have demonstrated its ability to infect other grasses [128–132]. Using qualitative and quantitative infection assays under greenhouse conditions, we could demonstrate a strong extent of host-specificity. Isolates from the *Aegilops* population 1 could only infect *Aegilops cylindrica* plants, while the isolates belonging to the *Aegilops* population 2 could only infect *Aegilops cylindrica* and *Aegilops tauschii*. The isolates from the wheat-infecting population could only infect *Triticum aestivum* and other closely-related *Triticum* species (e.g. *Triticum turgidum*), although we observed lesions in one *Aegilops tauschii* plant inoculated with a batch of wheat-infecting *Z. tritici* isolates. Considering that the *Z. tritici* populations analyzed in this study come from the Fertile Crescent region, and therefore from the most probable center of origin of the *Z. tritici* species, it is not surprising to find such host range among the isolates. Our infections assays and amplicon sequencing of herbarium specimens have pointed out that *Z. tritici* isolates infect closely-related grass species from the *Triticum*-*Aegilops* complex that participated in the domestication of common bread wheat (*Triticum aestivum*) in the Fertile Crescent [133–135]. Moreover, our demography analyses show that the divergence between *Aegilops*- and wheat-infecting *Z. tritici* populations occurred after 10,000 years ago, which coincides with previous findings of *Z. tritici* speciation event [36].

Due to the intrinsic properties of different plant species (e.g. resistance genes; antimicrobial metabolites), different host species can be considered as divergent habitats and thus lead to a strong selection on associated fungal pathogen populations [3,4,136]. A strong divergent selection due to host shift or host range expansion, if persistent over generations and coupled with a reduction of gene flow followed by reproductive isolation between populations, can drive these populations to a process of ecological speciation [3,6]. Regarding *Z. tritici*, it has been shown that divergence and reproductive isolation between *Zymoseptoria* species occurred with the presence of gene flow, which suggests that ecological divergence and continuous accumulation of minor incompatibilities may have contributed to the incipient speciation of this fungal species [19,36]. The complex evolutionary history with gene flow in different directions at different time points involving multiple *Zymoseptoria* species makes the implementation of introgression tests like ABBA-BABA or other F-statistics between *Z. tritici* populations challenging [40,137–139]. However, our analyses show that divergence between the *Z. tritici* populations on distinct hosts has been sufficiently strong to reduce the gene flow between these host-specific lineages, as evidenced by the lack of reticulate branches observed connecting the *Aegilops*-infecting *Z. tritici* isolates to the worldwide collection of wheat-infecting *Z. tritici*. Experimental crosses and further genealogical inferences using isolates from each host-specific lineage are required to determine if reproductive isolation, in pre- or post-zygotic stages, takes place and if the *Aegilops*-infecting *Z. tritici* can be considered a cryptic species.

## Conclusion

Altogether, this study provides evidences of distinct host-specific *Z. tritici* lineages and further suggests that the divergence observed between these lineages is part of an ongoing incipient speciation event. Using population genomics analyses coupled with infection assays, our results underline that host specificity is a strong determinant of the population structure and divergence of the *Z. tritici* species and support a host-tracking coevolutionary scenario. Our work also presents a valuable wild pathosystem (*Aegilops*-*Z. tritici*) that can be used for functional studies and further evolutionary analyses on the impact of domestication of plants and associated pathogens. At last, our findings highlight the importance of epidemiological surveillance in non-cultivated plant species considering their contribution as reservoirs and “green bridges” of multiple pathogen lineages [19,68], particularly in regions where wild and domesticated hosts are in close range and new diseases outbreaks are prone to emerge.

## Material and methods

### Fungal isolates and growth conditions

*Aegilops* spp. and wheat (*Triticum aestivum* L.) leaves with *Zymoseptoria* symptoms were collected at different locations in Iran (S1 Table). Fungal specimens were isolated, stored in cryo-stocks at −80 C° and the species confirmed using sequencing of the *ITS* and *Beta-tubulin* loci following protocols and primers described previously [42]. All fungal isolates used in this study were propagated at 18 C° in YMS medium (4 g yeast extract, 4 g malt, 4 g sucrose per 1 L, 20 g agar per L for plates; [171]) for 4-5 days prior use. For DNA extraction, fungal cells were retrieved directly from −80 C° glycerol stocks, grown in liquid YMS medium for 5 days at 18 C° under shaking at 200 rpm, and harvested by centrifugation (3,500 rpm for 10 min).

### Fungal DNA extraction and whole-genome sequencing

High-quality genomic DNA extractions were performed for all 310 *Z. tritici* isolates following a CTAB DNA extraction protocol described previously [42]. Prior to whole-genome sequencing, isolates were clone-corrected by PCR using ISSR (Inter Simple-Sequence Repeat) markers [140–142]. The list of primers used is described in S12 Table. Isolates originated from the same leaf lesion presenting the same ISSR pattern were considered clones and only isolates with different ISSR patterns were selected for genome sequencing. A subset of 123 clone-corrected isolates (S1 Table) were then submitted for whole-genome sequencing using paired-end reads of 150bp on an Illumina HiSeq 3000 platform. Library preparations and sequencing were performed at the Max Planck Genome Center, Cologne, Germany (http://mpgc.mpipz.mpg.de).

### Identification of herbarium species

The Iranian *Z. tritici* collections were obtained from grasses which identity at the species level was not always clear. Therefore, we performed Sanger sequencing of plant barcode loci to determine the species present in our herbarium material. To this end, a combined PTB-phenol protocol was used to extract genomic DNA from the herbarium leaves [42,143,144]. Briefly, 10 mg of leaves from each herbarium sample were place in 2mL sterile cryotubes containing three sterile 2,8 mm ceramic beads (biolab products GmbH, Bebensee, Germany) and snap-frozen in liquid nitrogen. Samples were then moved to a Precellys tissue homogenizer (Precellys Evolution, Bertin instruments GmbH, Frankfurt, Germany) and the plant tissue was ground for two cycles of 30 seconds with 5500 RPM and at – 15°C. After grinding, 1.2 ml of PTB-Buffer (PTB-stock solution, 0,4mg/ml proteinase K, 50mM DTT, 2,5 mM PTB) were added to the powdered plant material and the sample was stored overnight at 37°C under constant rotation [143,144]. Samples were then centrifuged for 10 minutes at 16.000 x g and the supernatant was added to 800μL of 1:1 phenol/chloroform solution. The next steps were followed as described at the CTAB DNA extraction protocol of Fagundes et al. 2020 (step 10 onwards; [42]). PCR amplification of the plant nuclear region *ITS2* was performed using the primers described in S12 Table and submitted for Sanger sequencing at Eurofins Genomics (Ebersberg, Germany). Homology-based searches were performed for each sequenced amplicon using nucleotide BLAST (blastn) [43] optimized for a discontiguous megablast search against the National Center for Biotechnology Information (NCBI) nucleotide database (https://blast.ncbi.nlm.nih.gov/Blast.cgi). To observe phylogenetic relationships, multiple sequence alignments were performed using ClustalW (https://www.ebi.ac.uk/Tools/msa/clustalo/) [145] and genetic distances were calculated using MEGA 5.2.2 [146] according to the Kimura 2-Parameter (K2P) model. Positions with less than 95% site coverage were eliminated from the alignments. Neighbor-Joining (NJ) consensus dendrograms were then constructed with 1000 bootstrap replicates also using MEGA 5.2.2 [146]. Additional barcode loci sequences from *Aegilops* and *Triticum* species as well as from barley (*Hordeum vulgare*) and rye (*Secale cereale*) were obtained from the NCBI GenBank database (https://www.ncbi.nlm.nih.gov/genbank/; S13 Table).

### Read mapping and variant calling

Raw sequencing reads were first trimmed for adapters and sequencing quality. Raw reads were trimmed using Trimmomatic v 0.39 [147] using the following parameters: LEADING:20 SLIDING-WINDOW:4:30 AVGQUAL:30 MINLEN:50. Trimmed reads of all isolates were mapped to the *Z. tritici* reference genome IPO323 [121] using the short-read aligner bwa-mem v. 0.7.17 [148]. Conversion, sorting, merging, and indexing of alignment files were performed using SAMtools v. 1. 13 [149]. PCR duplicates and read groups were determined using Picard tools v. 2.26.2 (https://broadinstitute.github.io/picard/). Single nucleotide polymophism (SNP) calling and variant filtration were carried out using the Genome Analysis Toolkit (GATK) v. 4.1.4.1 [150]. The GATK HaplotypeCaller was first used on each isolate individually with the commands -ERC GVCF and -ploidy 1. Then, gVCFs were merged using the GATK CombineGVCFs program and joint variant calling was performed on the merged gvcf variant file using GATK GenotypeGVFs. Only SNPs were maintained on the joint variant call file (VCF) using the GATK SelectVariants tool with the command --select-type-to-include SNP. We performed hard-filtering on SNPs based on quality cut-offs and following the GATK Best Practices recommendations [151] using the GATK VariantFiltration and SelectVariants tools. First, we filtered low depth genotypes using the options --genotype-filter-expression “DP < 3”, --set-filtered-genotype-to-no-call and --genotype-filter-name “low_depth”. Then, the following filter conditions were applied: DP > 5000.0; QD < 20.0; MQ < 50.0; FS > 20.0; ReadPosRankSum, MQRankSum, and BaseQRankSum between −2 and 2. SNPs that failed to pass these criteria were subsequently removed from the VCF file using GATK SelectVariants with the options --exclude-non-variants, --remove-unused-alternates and --exclude-filtered. Genotyping accuracy of GATK was previously analyzed in a population genomic study of *Z. tritici* and shown to be highly congruent with other SNP callers [28]. We further excluded SNPs located on accessory chromosomes (i.e., chromosomes not present in all isolates) and retained only biallelic SNPs with a genotyping rate of ≥ 90% using VCFtools v 0.1.13 [152]. Based on the 1,357,300 remaining SNPs, we computed the pairwise genomic relatedness among the 123 isolates by constructing an Identity-By-State (IBS) similarity matrix using PLINK v. 1.07 [153]. Individuals that showed an IBS similarity > 0.9999 to any other were excluded for further analysis as they represent highly similar isolates and potentially clones. The filtered dataset of 1,355,994 biallelic SNPs across 85 individuals was used for the subsequent population genomics analyses.

### Linkage disequilibrium analysis

In order to assess the extent of linkage disequilibrium (LD) for the *Z. tritici* collections (*Aegilops*- and wheat-infecting), we estimated the LD decay for SNPs located on chromosome 1. To this end, we retained only SNPs with a minor allele frequency (MAF) of ≥ 5% in each individual collection using VCFtools v 0.1.13 [152]. We calculated the LD coefficient of correlation (r2) between all marker pairs up to a distance of 20 kb or up to 100 kb using PLINK v. 1.07 [153]. The analysis was performed individually for the *Aegilops*- and wheat-infecting populations to prevent biases caused by possible population structure or demography. The pairwise r2 values were then plotted over genomic distance using the package ‘ggplot2’ v. 3.3.2 [154] in R software v. 3.6.3 [155]. We also determined the approximate genomic distance required to reach 50% LD decay for each collection.

### Population structure and population genomics analyses

Different methods to estimate the population structure between the wheat- and *Aegilops*-infecting *Z. tritici* isolates were used. For these analyses, we retained SNPs with a MAF of ≥ 5% and further pruned the filtered SNP dataset to select genome-wide SNPs at equidistant intervals of 12 kb along the chromosomes using VCFtools v 0.1.13 [152]. This pruning step ensured the removal of SNPs in strong linkage disequilibrium based on our analysis and on previous reports of linkage disequilibrium decay in *Z. tritici* populations [69,156]. Based on the 2507 remaining SNPs, we performed a Principal Component Analysis (PCA) using the R Bioconductor package ‘SNPRelate’ v. 1.20.1 [157]. Population structure was further inferred using the individual ancestry coefficients calculated by the sNMF v. 2.0 algorithm implemented in the R package ‘LEA’ [50,51]. The *K* parameter was tested between 1 to 10 with 100 repetitions for each tested *K* value. The best run was chosen and plotted based on the lowest cross-entropy value across the 100 repetitions for each *K*. Finally, we calculated nucleotide diversity per site (π) [158] in 100 kb sliding windows using the LD-unpruned SNP dataset and a MAF > 5% for each *Z. tritici* collection individually. Genotypic diversity calculations were obtained using VCFtools v 0.1.13 [152] with the haploid mode fork provided by Julien Y. Dutheil (https://github.com/vcftools/vcftools/pull/69).

### Whole genome *de novo* assemblies, mating types and phylogenomics analyses

To understand the evolutionary relationship between the sequenced isolates, we analyzed draft genome assemblies of different *Zymoseptoria* species. In addition to the dataset generated here, we used previously published population genome Illumina sequencing data of*Z. tritici* isolates collected in Oregon (US), Switzerland, Australia and Israel [69], and from the closely related *Zymoseptoria* species *Z. ardabiliae*, *Z. pseudotritici* and *Z. brevis* ([45,46,68,186; S1 Table). Illumina paired-end reads of ≥ 100bp were trimmed for adapter and sequencing quality using Trimmomatic v 0.39 [147] with the following parameters: LEADING:20 TRAILING:20 SLIDINGWINDOW:5:20 MINLEN:50. To account for older Illumina sequencing libraries (paired-end reads of 75bp), Trimmomatic v 0.39 settings were modified as follow: LEADING:10 TRAILING:10 SLIDINGWINDOW:5:10 MINLEN:50. We generated *de novo* genome assemblies for each isolate with SPAdes v. 3.14.1 [160] using the following parameters: --careful and a kmer range (-k) of ‘21,33,55,77’.

In order to infer evolutionary relationships between *Zymoseptoria* isolates, two phylogenomic methods were used. A Jukes-Cantor distance matrix was constructed using the Illumina draft genomes and the Andi v. 0.12 software [161]. Then, a Neighbor-net network was produced using the generated distance matrix and the SplitsTree v.4 software [162,163] to detect evidence of phylogenetic reticulation. We also generated a maximum likelihood (ML) tree based on filtered genome-wide SNPs using the substitution model TVM+F+P+N9+G4 and 100 bootstraps in the PoMo software implemented in IQ-TREE 1.6.12 [46,47,164,165]. PoMo estimates population genetic parameters (mutation rates and fixation biases) using genetic variation between and within species and populations and accounts for phylogenetic tree discordances as ILS (incomplete lineage sorting) [46,47]. The software analyzes the provided genetic variation and automatically chooses the best substitution model for the data using ModelFinder (-m MFP option) [166]. The ML tree was visualized using the online tool iTOL v.6 [167]. The VCF files for this step were generated using GATK v. 4.1.4.1 [150] and the *Z. tritici* reference genome IPO323 [121] as described above. After low depth genotype filtering, SNPs were quality-filtered using the GATK VariantFiltration and SelectVariants tools based on the following filter conditions: DP > 8000.0; QD < 20.0; MQ < 50.0; FS > 20.0; ReadPosRankSum, MQRankSum, and BaseQRankSum between -2 and 2. Only biallelic SNPs present in core chromosomes and with no missing genotypes were retained using VCFtools v 0.1.13 [152]. Using this VCF file, we also estimated population differentiation by calculating all pairwise Weir and Cockerham’s Fst fixation indices among the *Z. tritici* populations [48]. For this calculation and to avoid SNPs in high linkage disequilibrium, we further pruned the filtered SNP dataset to select genome-wide SNPs at equidistant intervals of 12 kb along the chromosomes using VCFtools v 0.1.13 [152] as previously performed. Fst calculations were obtained using VCFtools v 0.1.13 [152] with the haploid mode fork provided by Julien Y. Dutheil (https://github.com/vcftools/vcftools/pull/69).

To analyze the distribution of MAT1-1 and MAT1-2 idiomorphs in the sequenced *Aegilops*- and wheat-infecting populations, we performed nucleotide BLAST (blastn) [43] searchers (e-value 1e−6, identity ≥ 80%) in the draft genome assemblies of each sequenced isolate using the *Z. tritici* MAT1-1 and MAT1-2 loci as query sequences [52]. The number of idiomorphs were counted for each *Z. tritici* population individually and chi-square tests were performed in R v. 3.6.3 [155].

### Selective sweep analyses

Selective sweep scans were based on the SNPs for which we could assign their ancestral states. In order to identify ancestral SNP alleles, we analyzed whole-genome population sequencing data of the two sister species of *Z. tritici*, *Z. ardabiliae* and *Z. pseudotritici*. The same *Z.pseudotritici* and *Z. ardabiliae* isolates used for the phylogenomics analyses were used in these steps (S1 Table). Illumina paired-end reads of 150bp were trimmed for adapter and sequencing quality using Trimmomatic v 0.39 [147] with the following parameters: LEADING:20 TRAILING:20 SLIDINGWINDOW:5:20 MINLEN:50. Some of the previous genome datasets were generated with shorter reads. Therefore, to account for older Illumina sequencing libraries (paired-end reads of 75bp), Trimmomatic v 0.39 settings were modified as follow: LEADING:10 TRAILING:10 SLIDINGWINDOW:5:10 MINLEN:50. Read mapping and GATK SNP calling procedures were the same as described for the *Z. tritici* populations above. We merged all gvcf variant files and performed a final joint variant calling to include all *Z. tritici, Z. pseudotritici* and *Z. ardabiliae* isolates. We filtered SNPs based on quality cut-offs as described above with the following filter conditions: DP > 5000.0; QD < 20.0; MQ < 50.0; FS > 20.0; ReadPosRankSum, MQRankSum, and BaseQRankSum between -2 and 2. For each of the *Z. tritici* sister species, we retained only biallelic SNPs present in core chromosomes with a genotyping rate >50% and with no interspecific polymorphism using VCFtools v 0.1.13 [152]. To determine the ancestral state of variable positions in the *Z. tritici* genomes, we used *Z. ardabiliae* and *Z. pseudotritici* as outgroups. A *Z. tritici* SNP allele was assigned as ancestral if the allele was fixed in both *Z. ardabiliae* and *Z. pseudotritici* species.

Signals of selection in selective sweep analyses can be erroneously detected due to confounding factors such as demography [23,57]. To decrease the number of false positives, we used two methods to detect selective sweeps that are robust to demography effects: the Composite Likelihood Ratio (CLR) method, implemented in SweeD [58] and the μ statistic, implemented in the RAiSD software [59]. RAiSD is a composite evaluation test that simultaneously evaluates hallmarks of selective sweep regions by accounting changes in the Site Frequency Spectrum (SFS), the levels of LD, and the amount of genetic diversity along chromosomes [59]. Both methods were run for each core chromosome and each *Z. tritici* population individually at grid points of 1kb using only the SNPs for which we could assign the ancestral state. To avoid erroneous CLR and μ scores due to low SNP density, we calculated genome-wide SNP density in 50kb non-overlapping windows for each population and retained only grid point scores contained in windows with at least 100 SNPs. The 99.5th percentile score distribution for each test and for each population was used as a threshold to identify outlier grid points. Selective sweep regions were then determined based on the previous linkage disequilibrium estimations (see above). For the *Aegilops* population 1, outlier grid points were merged if their distance was less than 12kb, whereas for the wheat-infecting population outlier grid points were merged if they were within a 5kb distance apart. To account for potential high linkage disequilibrium blocks around these identified selective sweep regions, we further extended these regions by adding 10kb to each end of *Aegilops* population 1 regions and 3kb for the wheat population ones. At last, we added 15kb to each end of the identified selective sweep regions in both populations to visualize the surrounding genomic locations.

### Analysis of genomic content in selective sweep regions

Gene and Transposable Element (TE) content in each selective sweep region was analyzed using the gene and TE models and functional annotations of the *Z. tritici* IPO323 reference genome published previously [62–64]. Population genomics statistics (Dxy, Fst, Tajima’s D and nucleotide diversity, *π*) were calculated using VCFtools v 0.1.13 [152] with the haploid mode fork provided by Julien Y. Dutheil (https://github.com/vcftools/vcftools/pull/69) and with the scripts provided by Simon Martin (https://github.com/simonhmartin/genomics_general). Calculations were performed in non-overlapping windows of 1kb (Dxy, Fst and *π*) or 5kb (Tajima’s D). Linkage disequilibrium across the selective sweep region was calculated using VCFtools v 0.1.13 [152] and PLINK v. 1.07 [153]. To this end, we used SNPs with a MAF of ≥ 1% for each population individually. Heatmap plots were created using the R package “LDheatmap” [168] and gene/TE models were drawn using the R package “gggenes” (https://cran.r-project.org/web/packages/gggenes/index.html). For the Gene Ontology (GO) enrichment analyses, we used the R package “topGO” [169] with the GO terms for each gene model published previously [63]. *p* values were calculated using Fischer’s exact test applying the topGO algorithm “weight01” considering GO term hierarchy. We reported significant GO categories for the “Biological Process” ontology when *p* ≤ 0.05. PFAM domain enrichment analyses were performed using a custom python script [56] with *p* values being calculated using chi-square tests. Lastly, we performed the McDonald & Kreitman (MK) test [170] to test if non-neutral evolution (e.g. positive selection) is acting on any of the genes in the selective sweep region. For this, we used the script “vcf2fasta.py” (https://github.com/santiagosnchez/vcf2fasta) to obtain gene sequences with polymorphisms for each host-diverging *Z. tritici* population individually. The orthologs genes in the *Z. ardabiliae* Za17 isolate [62] served as outgroup for these tests. For some of the analyzed genes, no orthologs could be identified in *Z. ardabiliae* Za17 isolate, and therefore no MK tests could be performed on them (Zt09_chr_7_00296, Zt09_chr_7_00298, Zt09_chr_7_00303 and Zt09_chr_7_00308). The MK tests were calculated with the web server MKT (http://mkt.uab.es/mkt/MKT.asp; [171]) using the settings “Standard MKT” and “divergence corrected by Jukes & Cantor”.

### Demography analyses

To access the evolutionary history of the sequenced *Z. tritici* isolates, we surveyed the demography of the *Aegilops*- and wheat-infecting *Z. tritici* populations using MSMC2 [53,54]. For these analyses, we used a genomic dataset of 45 wheat-infecting isolates and 28 *Aegilops*-infecting isolates from the *Aegilops* population 1. SNP calling was performed with bcftools v. 1.3.1 [149] according to the pipeline suggested in Schiffels & Wang, 2020 [54]. Assuming a diploid genome model, we obtained a diploid homozygous data set, thus phasing step was omitted [54]. Subsequently, only one genome index per VCF file was considered for generating msmc-input files. Gene coding regions were included due to the high gene density of the *Z. tritici* genome. MSMC2 was run 20 times, each with 8 haplotypes - four from the wheat- and four from the *Aegilops*-infecting population. From these 20 combinations, we computed both the effective population size and the relative cross-coalescence rate (RCCR). We assumed a mutation rate of 3.3e^−8^ per cell cycle [55] and 1 generation of sexual reproduction per year as in previous studies of *Zymoseptoria* evolution [38].

### Virulence assays

We assessed the host specificity of the sequenced isolates collected from wheat and *Aegilops* spp. using comparative infection assays. Independent qualitative and quantitative experiments were performed under greenhouse conditions [∼20°C (day)/∼12°C (night) and ∼70% humidity with a cycle of 16hr day/8hr night] using different wheat accessions and wild grass relatives as hosts (S4 Table). Seeds of wild wheat and *Aegilops* accessions were obtained from Dreschflegel Bio-Saatgut (Witzenhausen, Germany) and from NordGen (Lomma, Sweden). Seeds of the wheat cultivar Riband were kindly provided by Jason Rudd (Rothamsted Research, Harpenden, United Kingdom). Seeds of the wheat cultivar Chinese Spring were kindly provided by Bruce McDonald (ETH Zurich, Switzerland) and seeds of the cultivar Titlis were obtained from DSP AG (Delley, Switzerland). For all experiments, blastospore cultures of *Z. tritici* isolates were used for inoculum preparation as described previously [42]. To ensure all host species were in similar sizes at the time of inoculation, we used 14-days old seedlings of the wheat cultivars Chinese Spring, Riband and Titlis, and 20 days-old seedlings of all the other wild wheat and *Aegilops* accessions. Seed sowing and plant preparations were performed as described previously [42].

Qualitative disease scoring was performed manually at 21or 28 days post-inoculation (dpi) and inoculation procedures were done as previously described [42]. In order to avoid other biotic factors to be confounded with disease symptoms as e.g. senescence, isolates were only considered to be virulent on a specific host if at least one of the inoculated plants exhibited visual presence of pycnidia at 21 or 28 dpi. In the first experiment, isolates were inoculated in “batches” of populations containing 12 isolates each, namely “*Aegilops* pop 1” batch, representing the *Aegilops* population 1, “*Aegilops* pop 2” batch, representing the *Aegilops* population 2 and the “Wheat pop” batch, representing the wheat population. Isolates were selected to represent all herbarium samples and lesions of origin and to have an equal number of isolates per batch. In addition, we selected isolates based on the IBS (identity-by-state) scores to ensure genetically different isolates in the three “batches”. The most genetically differentiated isolates showing the lowest IBS score per herbarium sample and per lesion were chosen. We also included a “mock” treatment comprising sterile water with 0.1% Tween 20 (Roth, Karlsruhe, Germany), as well as a “positive” control, the reference *Z. tritici* isolate IPO323 (strain Zt244) known to be virulent on the wheat cultivar Riband. Blastospore suspensions for each isolate were individually adjusted to an OD600nm = 0.2 (approximately 1×10^6^ cells/mL) in 0.1% (v/v) Tween20 (Roth, Karlsruhe, Germany) when the different isolates were merged into the batches. Each batch was inoculated on three plants of each of the 31 plant hosts described in S4 Table. For a second qualitative experiment, we confirmed the observed virulence phenotypes of the *Aegilops* population 1 by individually inoculating the *Z. tritici* isolates on three plants of *Aegilops cylindrica* and of the wheat cultivar Riband. We selected two to three sequenced isolates per herbarium sample and per lesion based on the lowest IBS (identity-by-state) scores and adjusted the inoculum concentrations as in experiment one (OD600nm = 0.2). The mock treatment and the reference *Z. tritici* isolate IPO323 (strain Zt244) were used as control treatments.

At last, to get insights of the infection progress and to quantify virulence in the newly described *A. cylindrica*-*Z. tritici* pathosystem, we performed a disease progression and quantitative virulence analyses in a subset of isolates from the *Aegilops* population 1. For the disease progression analyses, one sequenced isolate per herbarium sample was selected based on the symptoms observed in the qualitative virulence assays. Inoculum concentrations were adjusted to 1×10^5^ cells/mL in 0.1% (v/v) Tween20 (Roth, Karlsruhe, Germany) and each *Aegilops*-infecting isolate was inoculated on 40 *A. cylindrica* plants following the same inoculation procedures described previously [42]. To compare the disease progression to a wheat-infecting isolate, we also inoculated 40 wheat cultivar Riband plants with an adjusted blastospore suspension (1×10^5^ cells/mL) of the *Z. tritici* reference isolate IPO323 (strain Zt244). Each inoculated leaf was manually inspected between 7 and 21 dpi and the occurrence of first visible symptoms (necrosis and pycnidia) recorded every two days. Mock treatments were used as a negative control. For the quantitative virulence assay, we used the same *Z. tritici* isolates and inoculum concentrations used for the disease progression analyses. *Aegilops*-infecting *Z. tritici* isolates were inoculated on to 50 *A. cylindrica* plants and the *Z. tritici* reference isolate IPO323 (strain Zt244) was inoculated on 50 wheat plants (cultivar Riband). Mock treatments were inoculated on 37 plants of each species as negative controls. At 21 dpi, individual inoculated leaves were scanned and analyzed using automated image analysis as previously described [42]. Pycnidia density (number of pycnidia per square centimeter of leaf surface) was used as readout of virulence. Statistical analyses were conducted in R version 4.0.5 [155].

### Confocal microscopy analyses

In order to further characterize the infection progress of *Aegilops cylindrica*-infecting *Z. tritici* isolates, we analyzed the morphological development of the *Z. tritici* isolate Zt469 on *A. cylindrica* plants using confocal laser scanning microscopy (CLSM). The *Z. tritici* isolate Zt469 was part of the *Aegilops* population 1 dataset for both population genomics and virulence assays and it was randomly selected to represent this host-specific population. *Aegilops cylindrica* plants were inoculated with Zt469 blastospore using an adjusted suspension of 1×10^5^ cells/mL in 0.1% (v/v) Tween20 (Roth, Karlsruhe, Germany) as described previously [42]. Mock treatment (sterile water with 0.1% Tween 20) was used as a negative control. Plants preparation and inoculation procedures were as described in the “Virulence assays” section above. We harvested inoculated *A. cylindrica* leaves at 7, 10, 12, 15, 18 and 21 days post-inoculation (dpi) to investigate the compatible interaction between Zt469 and the wild grass host. These time points were chosen to cover the expected four infection stages of *Z. tritici* ranging from early biotrophy to late necrotrophy as previously described [56]. Mock-treated plants were also harvested as negative controls. Leaf material was prepared and stained with wheat germ agglutinin (WGA) conjugated to fluorescein isothiocyanate (WGA-FITC) in combination with propidium iodide (PI) as performed previously [56]. In total, we examined 36 *A. cylindrica* leaves for Zt469, 8 leaves for mock treatment and analyzed up to 43 infection events per leaf sample by CLSM, creating a total of 44 confocal image z-stacks. Microscopy was conducted using a Zeiss LSM880 microscope (Carl Zeiss Microscopy, Germany). FITC was excited at 488 nm (argon laser) and detected between 500 and 540 nm. PI was excited at 561 nm (diode-pumped solid-state laser) and detected between 600 and 670 nm. We used the ZEN black and ZEN blue (Carl Zeiss Microscopy, Germany) software for analyses, visualization, and processing of image z-stacks. Animations of image z-stacks in “.avi” format can be played in VLC media player (available at http://www.videolan.org/vlc/) and are available from [172].

The infection stages were determined by examining central leaf sections (1-2cm) of the harvested leaves by CSLM as described previously [56]. Considering the heterogeneity of infection events happening in a single leaf [173], we determined the infection stage of each leaf sample by observing the most progressed infection events (ranging from stage “A” to “D” as previously described by [56]) along a transect of six 0.25 mm2 squares of leaf area. That is, if in a leaf transect, we observed the infection events representing specific infection stages as e.g. stage “B” and stage “C”, we assigned the infection stage “C” to the respective leaf sample. The determination and assignment of infection stages were performed in a single-blinded procedure in which the “examiner” did not know the time point post-inoculation of the leaf sample *a priori*.

### Transcriptome sequencing

We generated infection stage-specific transcriptome data based on *A. cylindrica* leaf material collected and analyzed by confocal laser scanning microscopy (CSLM) during infection with the *Z. tritici* isolate Zt469. Tissue from the same inoculated leaf was used for both RNA isolation and microscopy analyses. Four infection stages were determined ranging from early biotrophy [stage “A”; 7days post-inoculation(dpi)] to late necrotrophy (stage “D”; 21 dpi). Per infection stage, we selected the three most representative samples of that time point as biological replicates for total RNA extraction and subsequent transcriptome sequencing. Each biological replicate consisted of material from three inoculated *A. cylindrica* leaves that were pooled and homogenized in liquid nitrogen. We isolated total RNA from Zt469-infected and mock-treated *A. cylindrica* leaves using the Direct-zol RNA MiniPrep kit (Zymo Research, Irvine, CA, US) following the manufacturer’s instructions. Preparation of strand-specific RNA-seq libraries including polyA enrichment was performed at the Max Planck Genome Center (MPGC), Cologne, Germany (http://mpgc.mpipz.mpg.de). Library sequencing was also performed at the MPGC on an Illumina HiSeq3000 platform yielding paired-ends reads of 150 nt.

For RNA-seq of Zt469 during *in vitro* growth, we used the same dataset generated and analyzed in (Fagundes et al. 2024) consisting of three biological replicates. Briefly, Zt469 fungal spores were grown at 18 C° on YMS plates for three to four days and total RNA extracted using TRIzol (Invitrogen, Karlsruhe, Germany) following the manufacturer’s instructions. Preparation of RNA-seq libraries and sequencing was performed by Admera Health (South Plainfield, NJ, USA) on an Illumina HiSeq 3000 platform obtaining paired-end reads of 150 nt.

For the transcriptome data of the IPO323 isolate, we reanalyze previously generated RNA-seq datasets during *in vitro* growth on YMS medium and during infection at four different stages in the wheat cultivar Obelisk [56,174]. Infection stages also ranged from early biotrophy (stage “A”) to late necrotrophy (stage “D”) and a total of two biological replicates for the *in vitro* growth and per infection stage were analyzed.

### RNA-seq read mapping and gene expression analyses

We compared the expression of the genes localized in the selective sweep region on chromosome 7 in IPO323 and Zt469 genomes during *in vitro* growth and *in planta* infection. Raw sequencing reads were first trimmed using Trimmomatic v 0.39 [147] and low-quality nucleotides (Q<20) were further masked using FASTX-toolkit v0.0.13 (http://hannonlab.cshl.edu/fastx_toolkit/). Reads were then aligned to the Zt469 genome assembly (Fagundes et al. 2024) and to the reference genome of IPO323 [121] using HISAT2 [175]. Conversion, sorting, merging and indexing of alignment files were performed using SAMtools v. 1.7 [149]. For the estimate of gene expression, we calculated the relative abundance of gene transcripts in FPKM (Fragments per Kilobase of Transcript per Million fragments mapped) using the Cuffnorm software from the Cufflinks package v.2.2.1 [176]. Homologous candidate effector genes in Zt469 were identified by mapping the gene models of IPO323 on the Zt469 genome assembly (Fagundes et al. 2024) using Geneious v.2020.1.2 software (https://www.geneious.com/home/).

### Bacterial strains and culturing conditions

For the transformation process of fungal cells, plasmids were maintained in *Escherichia coli* TOP10 cells (Invitrogen, Karlsruhe, Germany) and the *Agrobacterium tumefaciens* strain AGL1 was used for *Agrobacterium tumefaciens*-mediated transformation (ATMT). Both bacterial strains were cultivated on dYT medium (double-yeast-tryptone, 1.6% [w/v] tryptone, 1% [w/v] yeast extract, 0.5% [w/v] NaCl and 2% Bacto agar for solid medium) and grown at 28°C and 37°C for *A. tumefaciens* and *E. coli*, respectively. For plasmid selection, *E. coli* cells were grown in dYT medium supplemented with 40 μg/mL kanamycin (Sigma, Taufkirchen, Germany) and at 200 rpm for liquid cultures. For the *A. tumefaciens* strain AGL1, we grew the cells in dYT medium containing 50 μg/ml Rifampicin (Sigma, Taufkirchen Germany) and 100 μg/ml Carbenicillin (Sigma, Taufkirchen, Germany) to maintain the plasmids already present in the AGL1 strain, and 40 μg/mL kanamycin (Sigma, Taufkirchen, Germany) were added for plasmid selection.

### Generation of plasmid and ATMT of *Z. tritici* cells

To access the functional role of the candidate effector gene Zt09_chr_7_00299 in the *Aegilops*-infecting *Z. tritici* isolate Zt495, we generated targeted gene deletion mutant strains using *Agrobacterium tumefaciens*-mediated transformations (ATMT) as described previously [174,177,178]. Plasmid construct and overlapping primer pairs were designed using Geneious v.2020.1.2 software (https://www.geneious.com/home/; S12 Table). The plasmid pES61, a derivate of the binary vector pNOV-ABCD [177] which contains a kanamycin resistance for selection of *E. coli* transformants, was used as backbone vector for the plasmid construct. For targeted gene deletion, we amplified two DNA fragments of 1 kb upstream and downstream of the candidate effector gene Zt09_chr_7_00299 on Zt495 by PCR. These fragments served as flaking regions to facilitate homologous recombination of the selective hygromycin resistance marker cassette (*hph*; [177]) with the recipient Zt495 genome at the correct genomic location. Genome coordinates of the genes in the selective sweep region in Zt495 were annotated by mapping the gene models of IPO323 against the Zt495 draft genome assembly using nucleotide BLAST in Geneious v.2020.1.2 software (https://www.geneious.com/home/). We assembled the DNA fragments up- and downstream of the Zt09_chr_7_00299 ORF with the resistance marker cassette and ligated them into the restriction enzyme-digested plasmid pES61 using Gibson assembly [179].

For ATMT of *Z. tritici*, the assembled plasmid was amplified in *E. coli* TOP10 cells and transformed in electro-competent *A. tumefaciens* AGL1 cells using standard procedures. The plasmid construct was then introduced into Zt495 following an ATMT protocol described previously [67]. Transformed Zt495 colonies were visible on YMS plates supplemented with 150 μg/mL Hygromycin (Roth, Germany) two weeks after the ATMT procedure. Single-cell colonies were then propagated in YMS medium for further DNA extraction and confirmation of correct homologous recombination by PCR followed by Southern blot analyses [180]. For confirmation of transformed strains using PCRs, we amplified the resistant marker cassette and flanking regions using the outer primers of the deletion constructs. To generate the Southern blot probes, we amplified the resistance marker cassette and the downstream flank of the deletion construct using the PCR Digoxigenin (DIG) labeling mix (Roche, Mannheim, Germany) following manufacturer’s instructions. List of all primers designed and used for these steps can be found at S12 Table.

### Isolation of fungal DNA for PCR screening and Southern blot analyses

For confirmation of transformed strains using PCRs, we extracted DNA from single *Z. tritici* colonies using a rapid genomic DNA extraction protocol described previously [42]. For Southern blot analyses, we extracted DNA from five-days old *Z. tritici* blastospore cultures using a standard phenol-chloroform extraction protocol [181].

### Virulence assays of *Z. tritici* deletion mutants

To assess potential effects of the candidate effector Zt09_chr_7_00299 gene deletion on pathogenicity, we performed a quantitative virulence assay under greenhouse conditions (16/8 hours light/dark, ∼20/10°C day/night and ∼70% rel. humidity) using *Aegilops cylindrica* as a host. The procedures for plants preparation and *Z. tritici* blastospore inoculation were performed as described in the “Virulence assays” section. We inoculated *A. cylindrica* plants with adjusted blastospore suspensions of 1×10^6^ cells/mL in 0.1% (v/v) Tween20 (Roth, Karlsruhe, Germany) of the Zt495 wild-type and the three deletion mutant strains. Thirty plants were inoculated per strain. Mock treatment (sterile water with 0.1% Tween 20) was used as a negative control and also inoculated on thirty plants. Leaves were harvested at 21 dpi, scanned, and analyzed using automated image analysis as previously described [42]. We used pycnidia density (number of pycnidia per square centimeter of lesion) as a readout of virulence. Statistical analyses were conducted in R version 4.0.5 [155]. Outlier values were removed when the observed value was 1.5 times the interquartile range more than the third quartile (Q3) or 1.5 times the interquartile range less than the first quartile (Q1).

## Supporting information

Fagundes_2024_Zymo_PopGen_Supplementary_Material

## Data availability

The datasets generated and analyzed during the current study are openly available in different repositories. Whole-genome sequencing raw reads (FASTQ files) are available in the NCBI Short Read Archive (SRA; https://www.ncbi.nlm.nih.gov/bioproject/?term=) under the BioProject Accession Number: PRJNA1162695. *In planta* and *in vitro* RNA-seq datasets for Zt469 are available at the NCBI SRA BioProject accession number PRJNA1162778. Long-read genome assembly of the Zt469 isolate as well as Zt469 gene annotation from Fagundes et al. 2024 is available at https://doi.org/10.5281/zenodo.13773246. The genome sequence of the reference *Z. tritici* isolate IPO323 (MYCGR v2.0) is available at NCBI under the RefSeq assembly GCF_000219625.1.

## Acknowledgements

The authors would like to thank Danilo Pereira for support with the genomics analyses, Janine Müller for support with the infection assays, and all members of the Environmental Genomics group for helpful discussions.

## Supporting Information

**S1 Fig. Genome-wide distribution of IBS values per host collection.** Boxplot showing the distribution of IBS (Identity-By-State) values calculated genome-wide by pairs of *Z. tritici* isolates within each collection. P-value was calculated using Wilcoxon rank sum test.

**S2 Fig. Maximum-likelihood (ML) tree based on intraspecific *Zymoseptoria tritici* polymorphisms**. ML tree was constructed based on genome-wide biallelic SNPs with no missing genotypes across the different *Z. tritici* populations using the software PoMo [46,47] and 100 bootstraps.

**S3 Fig. Pairwise divergence values (Fst) between *Zymoseptoria tritici* populations**. Heatmap showing genome-wide mean divergence values based on the Weir & Cockerham’s Fst fixation indices [48]. Mean Fst values were calculated using biallelic SNPs with no missing genotypes across the different *Z. tritici* populations. Darker colors indicate higher divergence between pairs of populations.

**S4 Fig. Phylogenetic network between *Zymoseptoria* species**. Neighbor-net network based on whole genome distances. In the *Z. tritici* cluster, red ellipse represents the two *Aegilops*-infecting *Z. tritici* populations while the blue elliptical shape represents worldwide wheat-infecting *Z. tritici* populations. Geographical locations of origin are also indicated. Scale bar indicates branch distances.

**S5 Fig. Principal component analysis on genome-wide independent SNPs among host-diverging *Zymoseptoria tritici* isolates. (A)** PCA showing the second and third principal components. Percentage of variance explained by each component is shown in parentheses. **(B)** Lollipop plot showing the percentage of variance explained by the first 12 PCs.

**S6 Fig. Admixture analyses of host-diverging *Zymoseptoria tritici* isolates. (A)** Ancestry coefficients and clustering assignments using the sNMF v2.0 software [50,51]. Each vertical bar represents an isolate from the *Aegilops* (red arrow) and wheat-infecting (blue arrow) *Z. tritici* collection with each color indicating one genetic cluster. The color height in each vertical bar represents the probabilities of cluster assignment based on genome-wide independent SNPs. **(B)** Dot plot showing the cross-entropy values calculated for each ancestral population tested (K=1 to K=10). Values represent the lowest score over 100 independent runs for each K.

**S7 Fig. Herbarium species correlate with *Zymoseptoria tritici* population structure.** Neighbor-Joining (NJ) dendrogram performed with 1000 bootstrap replicates based on multiple sequence alignment of plant ITS2. The genetic distances were calculated according to the Kimura 2-Parameter model. Numbers at the branches show bootstrap values. Sequences amplified from greenhouse-grown plants (S4 Table) are indicated. Additional sequences were obtained from the NCBI database and are marked with asterisks and accession number. The *ITS2* sequence of *Hordeum vulgare* was used as an outgroup. List of accession numbers and references are described in S13 Table.

**S8 Fig. Nucleotide diversity and linkage disequilibrium decay in host-diverging *Zymoseptoria tritici* populations. (A)** Distribution of genome-wide nucleotide diversity (*π*) values calculated for each *Z. tritici* population individually. P-values were calculated using pairwise Wilcoxon rank sum tests. **(B)** Linkage disequilibrium decay across 100 kb of chromosome 1. Top, brown line represents the LD decay for *Aegilops*-population 2; mid, red line represents the LD decay for the *Aegilops* population 1 while the bottom, blue line represents the LD decay for the wheat-infecting population. Dashed perpendicular lines represent distance to reach 50% maximum LD value in each population.

**S9 Fig. Effective population size (Ne) of the *Aegilops*- and wheat-infecting *Zymoseptoria tritici* populations.** The effective population size was calculated for each Iranian population individually using 20 different combinations of 4 haploid genotypes for the wheat-infecting population **(A)** and 20 different combinations of 4 haploid genotypes for the *Aegilops* population 1 **(B)**. Each line in both plots represents the different genotype combinations. Grey, vertical bars represent timeframe of wheat domestication and *Z. tritici* speciation (∼10.000 years ago; [36,133–135]). Time is indicated as thousands of years (Ky). Calculations were performed using MSMC2 [53,54] assuming a mutation rate of 3.3e-8 per cell cycle [55] and 1 generation of sexual reproduction per year [38].

**S10 Fig. Relative cross-coalescence rate (RCCR) between Iranian host-diverging *Zymoseptoria tritici* populations**. The RCCR was calculated using 20 different combinations of 8 haploid genomes, 4 from wheat-infecting and 4 from the *Aegilops*-infecting *Z. tritici* population 1. Each line represents a different combination of samples between the two populations. Grey, vertical bar represents timeframe of wheat domestication and *Z. tritici* speciation (∼10.000 years ago; [36,133–135]. Time is indicated as thousands of years (Ky). Calculations were performed using MSMC2 [53,54] assuming a mutation rate of 3.3e-8 per cell cycle [55] and 1 generation of sexual reproduction per year [38].

**S11 Fig. Summary of qualitative virulence assay performed by *Z. tritici* population batches**. In this experiment, batches of twelve *Z. tritici* isolates based on the identified Iranian populations, namely “*Aegilops* pop 1” batch, representing the *Aegilops* population 1, “*Aegilops* pop 2” batch, representing the *Aegilops* population 2 and the “Wheat pop” batch, representing the wheat population were inoculated individually along mock and the reference *Z. tritici* isolate IPO323 (strain Zt244) as control treatments in 31 wheat and wild grasses accessions (S4 Table). Pictures show inoculated leaves at 21 days post-inoculation (dpi) and represent a subset of the results obtained. Each grey column represents leaves inoculated with a different population batch. Complete description of qualitative virulence results can be found at S5 Table.

**S12 Fig. Disease progression analyses on host-diverging *Zymoseptoria tritici* isolates.** To analyze the infection progression, we used a subset of *Aegilops*-infecting *Z. tritici* isolates from *Aegilops* population 1 along the *Z. tritici* IPO323 isolate (strain Zt244) and mock treatments as controls, and inoculated them individually at similar inoculum concentrations on its correspondent hosts: *Aegilops cylindrica*, for the *Aegilops*-infecting *Z. tritici* isolates and *Triticum aestivum* cv. Riband, for the *Z. tritici* IPO323 (strain Zt244). First signs of necrosis (yellow bars) and pycnidia (brown bars) were recorded every two days. Number of plants showing symptoms (n) are indicated.

**S13 Fig. *Aegilops*-infecting *Zymoseptoria tritici* isolate Zt469 shows four developmental stages during *Aegilops cylindrica* infection**. Micrographs obtained by CSLM (maximum projections of confocal image z-stacks) showing the four infection stages (stages “A” to “D”) of *Z. tritici* inside *A. cylindrica* leaves. Leaves were collected at the indicated time points (dpi; days post-inoculation). Mock samples (0.1% Tween in sterile water) were used as controls. Nuclei and *A. cylindrica* cells are displayed in purple and fungal hyphae in green. At stage “D”, two pycnidia harboring pycnidiospores are shown occupying substomatal cavities (white arrows). Scale white bars represent 50 μm. Raw image files and animation of z-stack micrographs of fungal-infected leaves in each stage are available at [172].

**S14 Fig. Candidate effector genes in selective sweep show different levels of expression in Zt469 and IPO323 *Z. tritici* isolates during host infection.** Normalized abundance of transcripts in FPKM for the three candidate effector genes in the selective sweep region of chromosome 7 in the reference wheat-infecting isolate IPO323 (top row) and their homologs in the *Aegilops*-infecting isolate Zt469 (bottom row). Relative expression of genes was measured during *in vitro* growth (in YMS media) and at four infection stages from early biotrophy (stage A) to late necrotrophy (stage D) in *A. cylindrica* (for Zt469) and wheat (for IPO323).

**S1 Table. *Zymoseptoria* isolates used in this study**. List of *Zymoseptoria* isolates used in this study indicating their geographical origin, year of collection, host and references.

**S2 Table. Nucleotide BLAST results of *ITS2* sequences from the herbarium specimens.** Homology-based searches were performed using nucleotide BLAST (blastn) [43] optimized for discontiguous megablast against the National Center for Biotechnology Information (NCBI) nucleotide database (https://blast.ncbi.nlm.nih.gov/Blast.cgi). Only the first three hits ordered by their percent identity values are listed for each sample.

**S3 Table. Mating type distribution between *Zymoseptoria tritici* populations.** Mating type idiomorphs were detected in all analyzed Iranian *Z. tritici* isolates using BLAST searches of both mating type loci (MAT1-1 and MAT1-2; [52]) sequences against whole-genome assemblies. Chi-square (*χ*2) tests were performed within each population based on the number of idiomorphs identified.

**S4 Table. List of host specimens used for the qualitative and quantitative infections assays under greenhouse conditions.**

**S5 Table. Qualitative results of greenhouse infection assays by *Zymoseptoria tritici* population batches.** Infection assays under greenhouse conditions were performed using 31 wild grass and wheat relatives (S4 Table) as hosts. In this experiment, *Z. tritici* isolates were inoculated by batches of 12 isolates each representing the three populations identified in this study: “*Aegilops* pop 1”, representing the *Aegilops* population 1; “*Aegilops* pop 2”, representing the *Aegilops* population 2; and the “Wheat pop”, representing the wheat population. Mock (0.1% Tween20 in sterile water) and the *Z. tritici* isolate IPO323 (strain Zt244) were used as controls. Hosts were considered susceptible to a specific population if at least one inoculated plant showed presence of pycnidia. Numbers in brackets represent proportion of plants with presence of pycnidia at 21 or 28 dpi over total number of plants inoculated. “+” represents compatible interactions (visual presence of pycnidia) and “-” represents incompatible interactions. Compatible interactions are highlighted in bold.

**S6 Table. Qualitative results of greenhouse infection assays by individual *Zymoseptoria tritici* isolates.** Infection assays were performed under greenhouse conditions. In this experiment, *Z. tritici* isolates were inoculated individually in *Aegilops cylindrica* and *Triticum aestivum* cultivar Riband plants (S4 Table). Mock (0.1% Tween20 in sterile water) and the *Z. tritici* isolate IPO323 (strain Zt244) were used as controls. Hosts were considered susceptible to a specific isolate if at least one inoculated plant showed presence of pycnidia. Numbers in brackets represent proportion of plants with presence of pycnidia over total number of plants inoculated. “+” represents compatible interactions (visual presence of pycnidia) and “-” represents incompatible interactions. Compatible interactions are highlighted in bold.

**S7 Table. Genomic coordinates of selective sweep regions identified in the two host-diverging *Zymoseptoria tritici* populations *Aegilops* population 1 and wheat population using the CLR and μ statistics methods.** Grey-colored cells represent overlapping regions commonly detected by the two selective sweep methods within each population. Red-colored letters represent regions commonly detected between the two populations.

**S8 Table. Summary metrics of selective sweep regions detected in the two host-diverging *Zymoseptoria tritici* populations *Aegilops* population 1 and wheat population using the CLR and μ statistics methods.**

**S9 Table. Summary of gene and TE models found in the selective sweep regions of *Aegilops* population 1 and wheat population using the CLR and μ statistics methods.** Gene/TE models and functional annotation of genes in the *Z. tritici* IPO323 reference genome were retrieved from previous studies [62–64].

**S10 Table. Gene ontology (GO) terms linked to biological processes significantly enriched in the selective sweep regions of the *Aegilops* population 1 and wheat population**. Only GO terms with P-value ≤ 0.05 are shown. P-values were calculated using Fischer’s exact test applying the topGO [169] algorithm “weight01” considering GO term hierarchy.

**S11 Table. PFAM domains significantly enriched in the selective sweep regions of the *Aegilops* population 1 and wheat population.** Only PFAM domains with P-value ≤ 0.05 are shown. P-values were calculated using a chi-square (χ2) test and a custom python script published previously [56].

**S12 Table. List of primers used in this study.**

**S13 Table. Specimens and *ITS2* GenBank accession numbers of the *Triticeae* species used for the herbarium phylogenetic analyses.**

