## Supplementary figures and images for "Host specialization defines the emergence of new fungal plant pathogen populations"

### S1_Fig.tiff

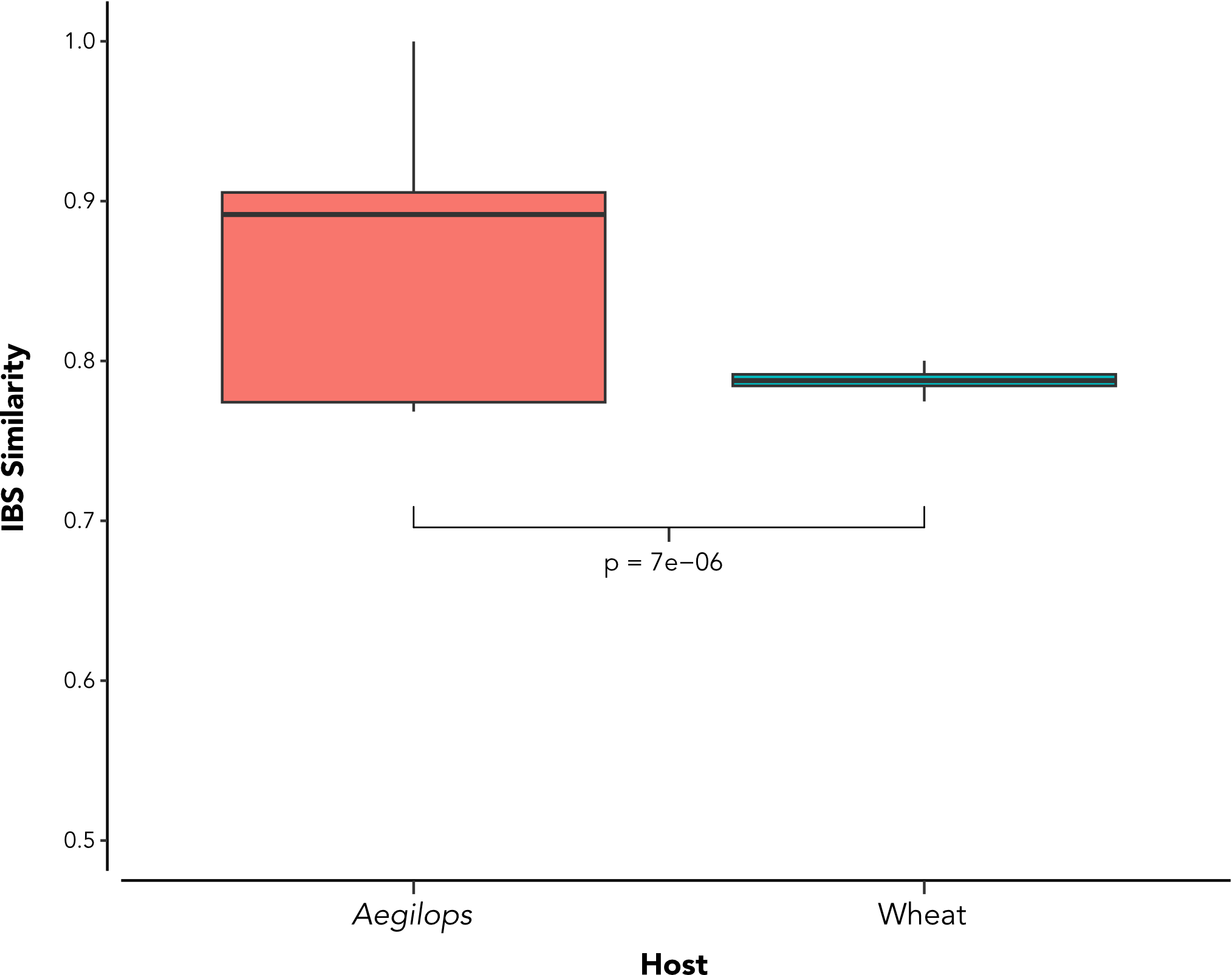

### S2_Fig.tiff

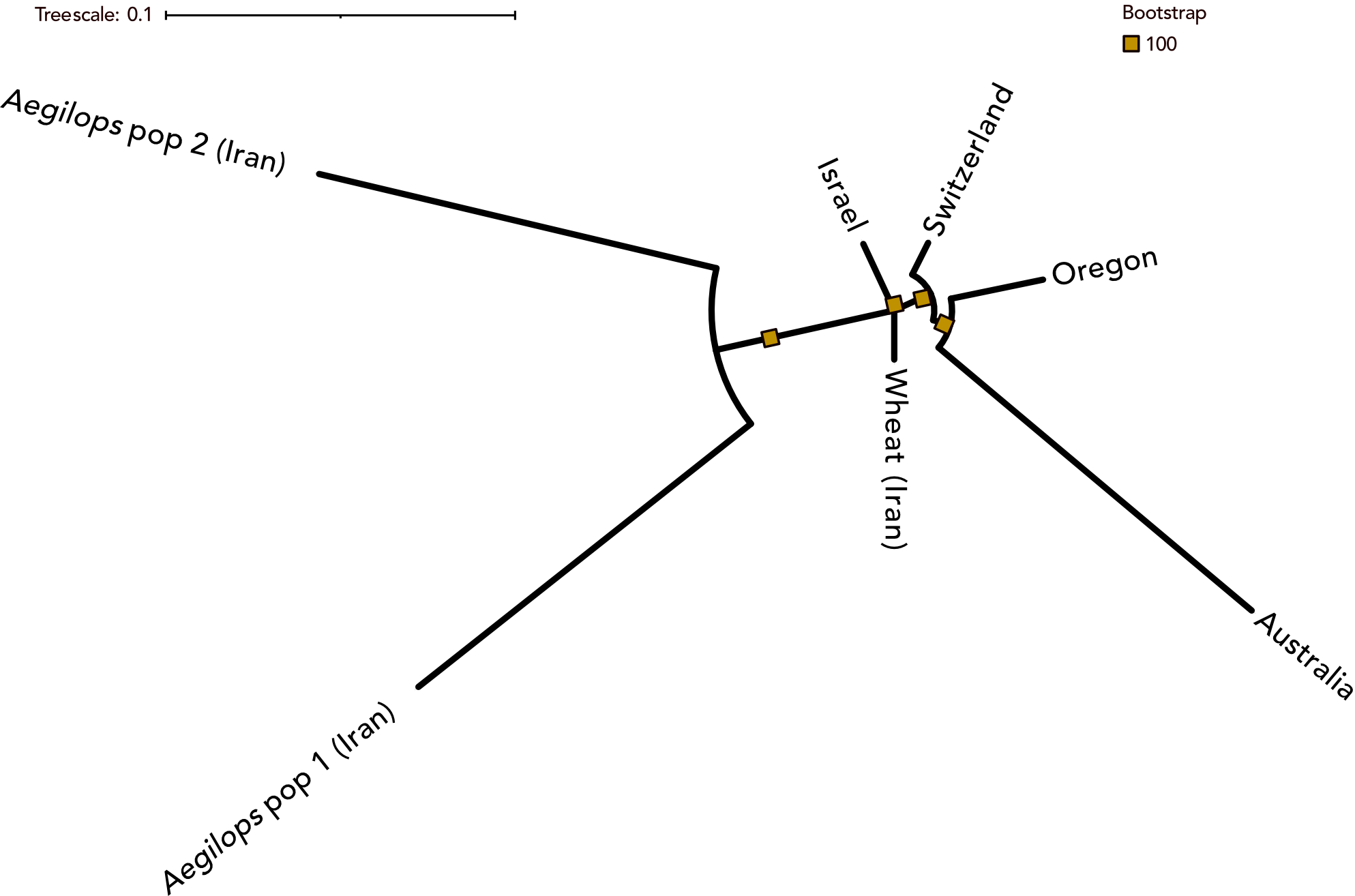

### S3_Fig.tiff

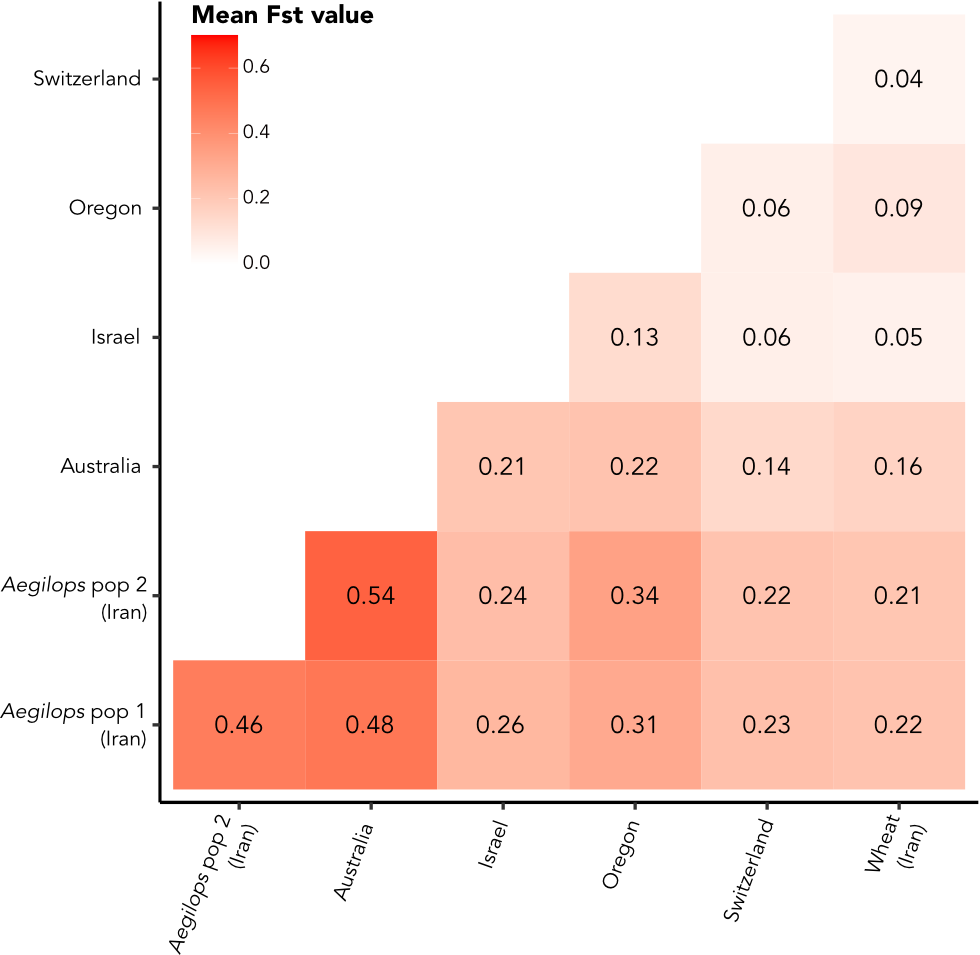

### S4_Fig.tiff

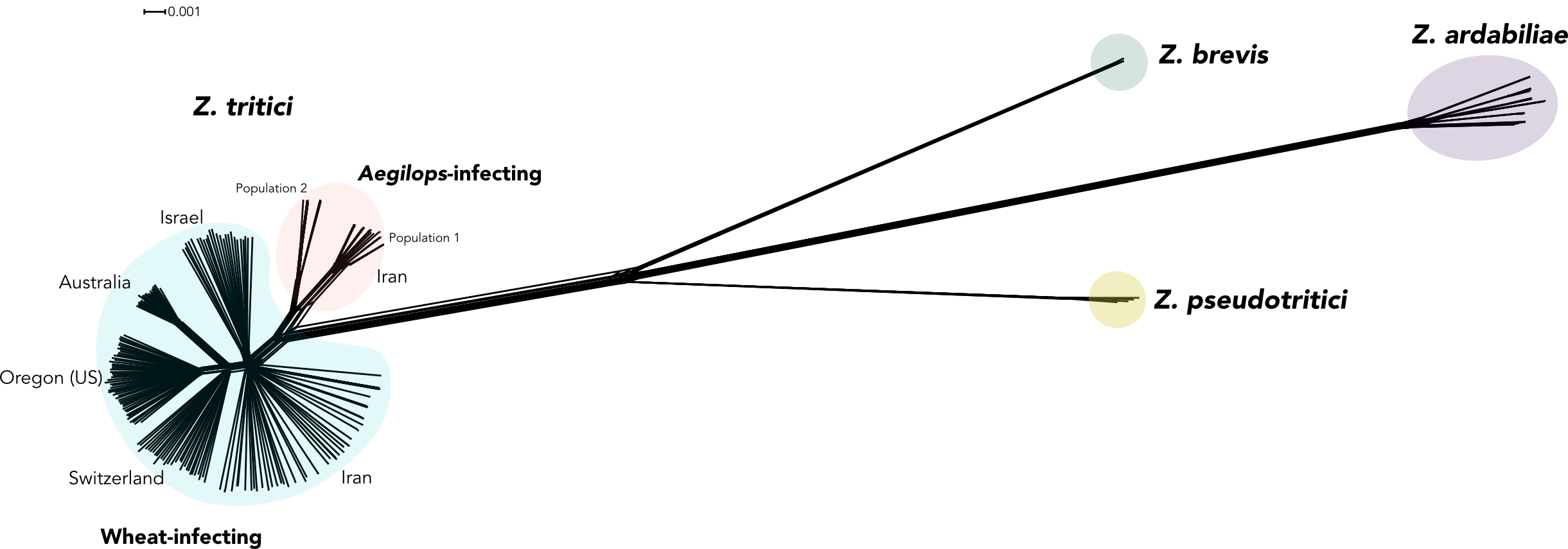

### S5_Fig.tiff

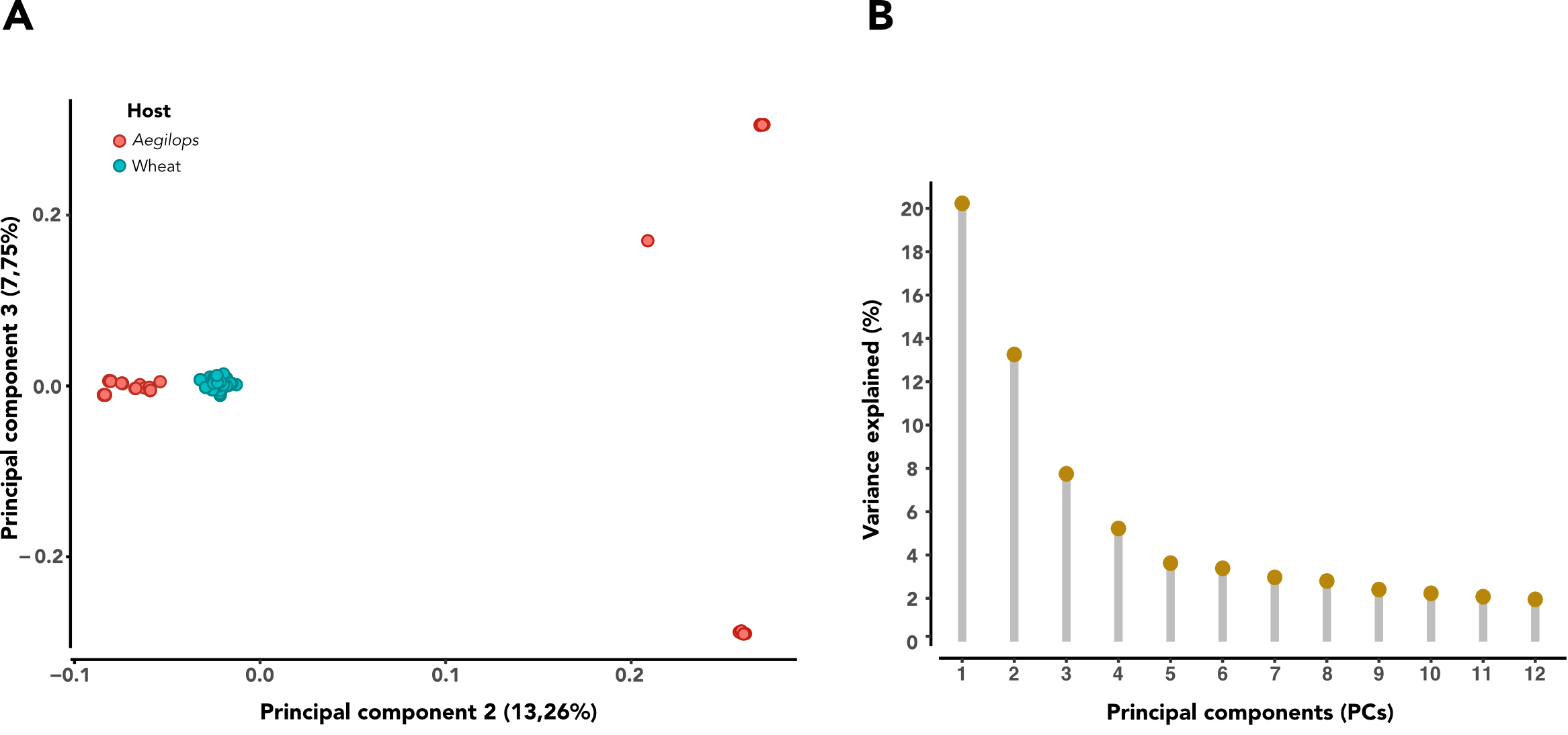

### S6_Fig.tiff

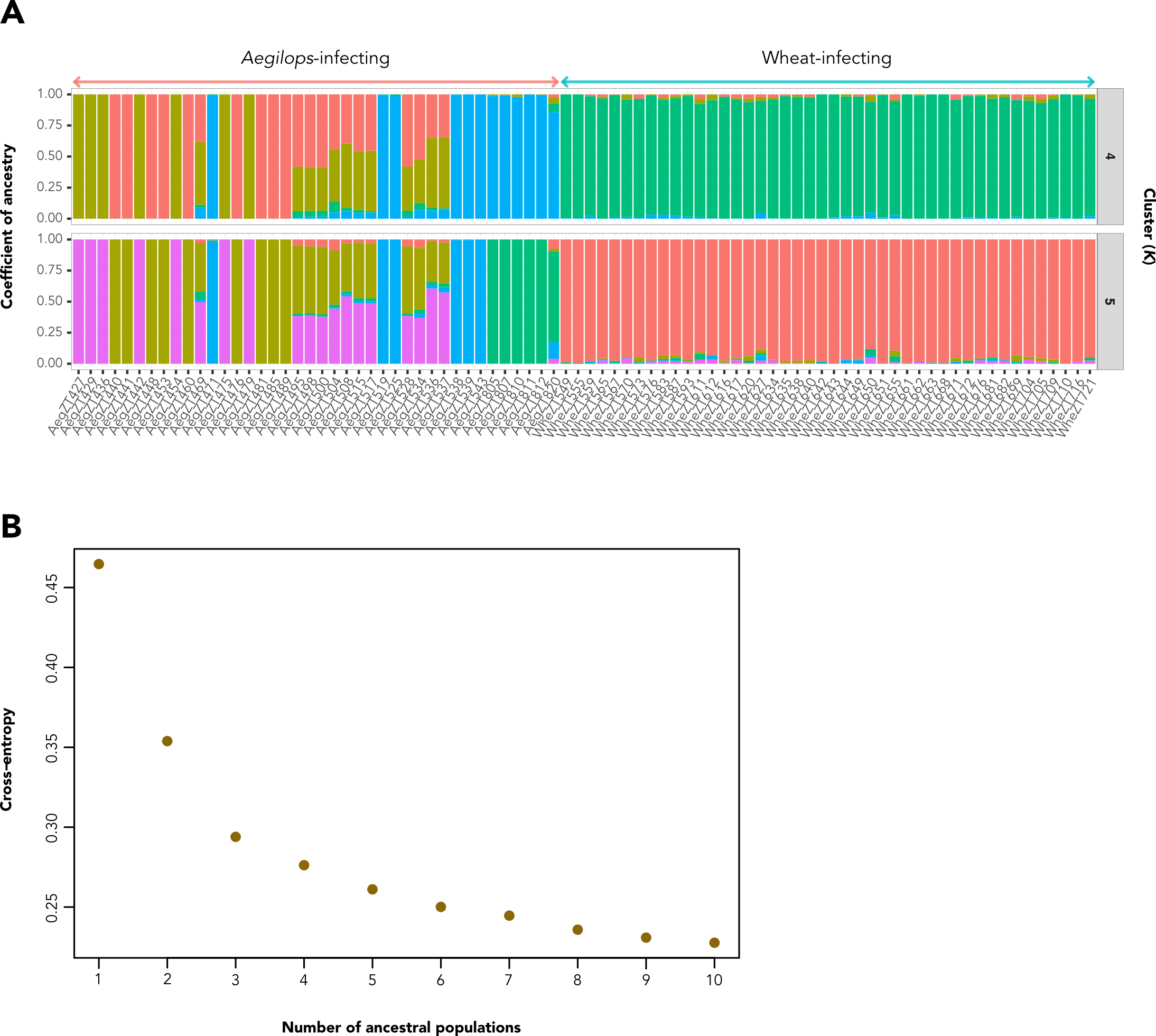

### S7_Fig.tiff

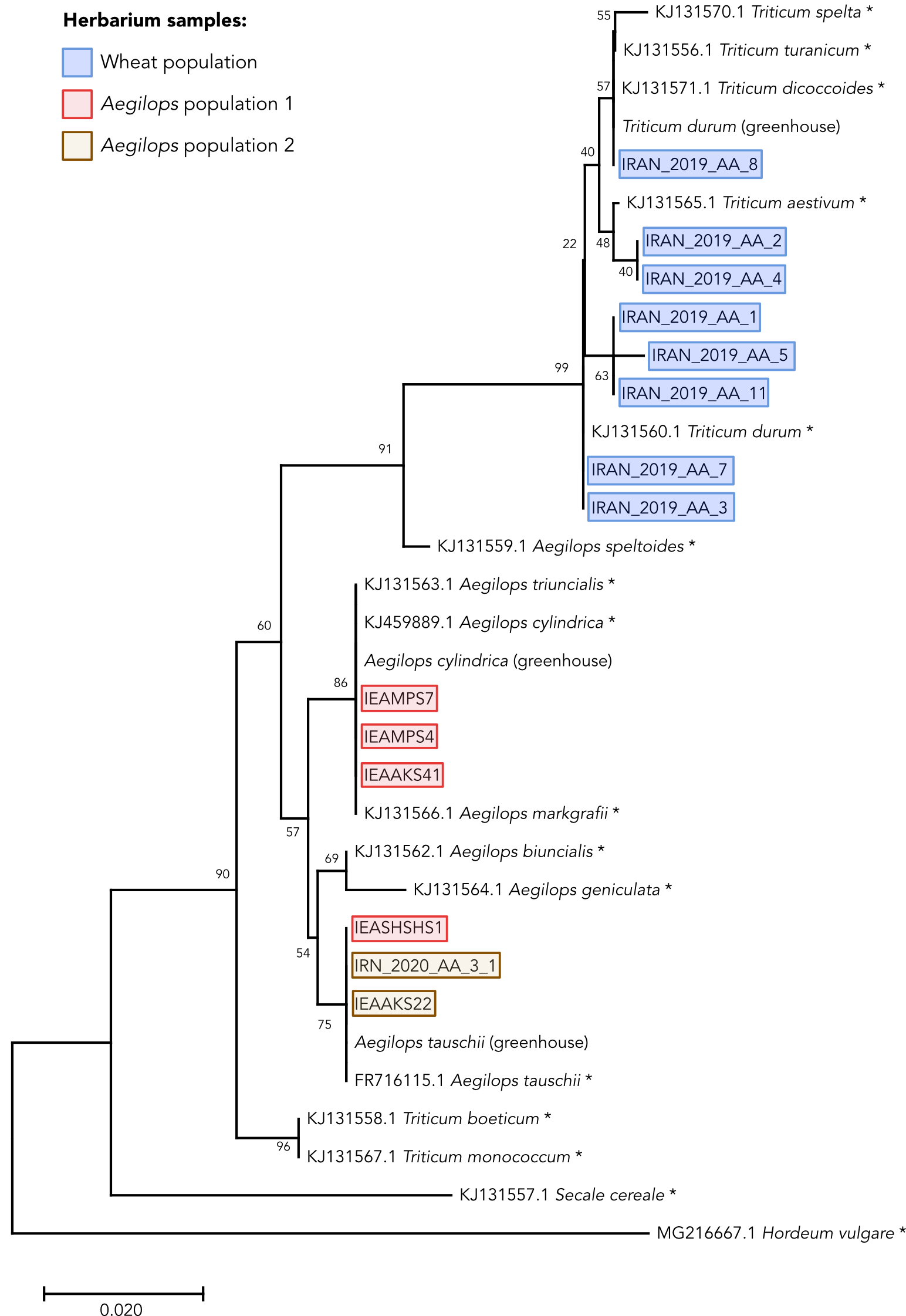

### S8_Fig.tiff

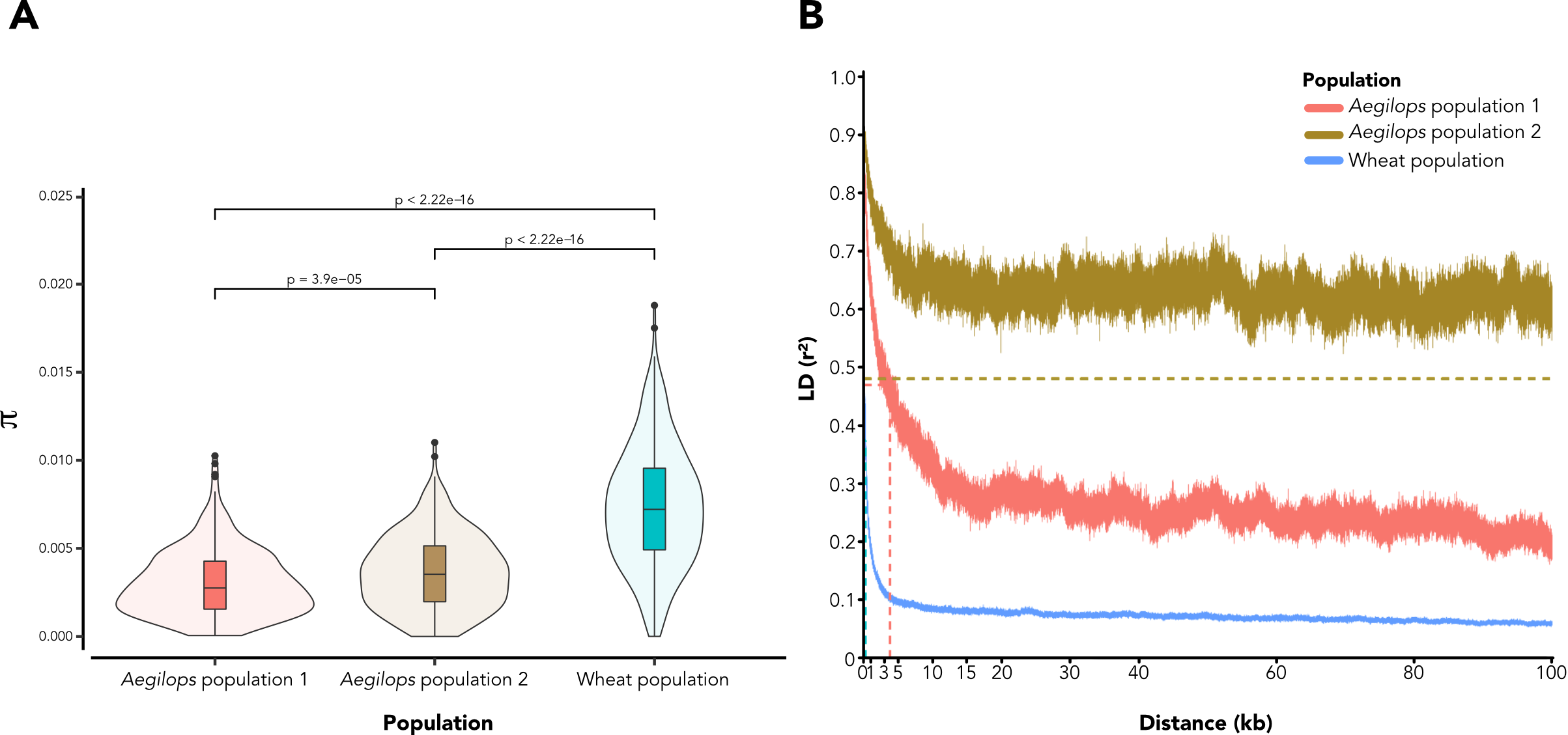

### S9_Fig.tiff

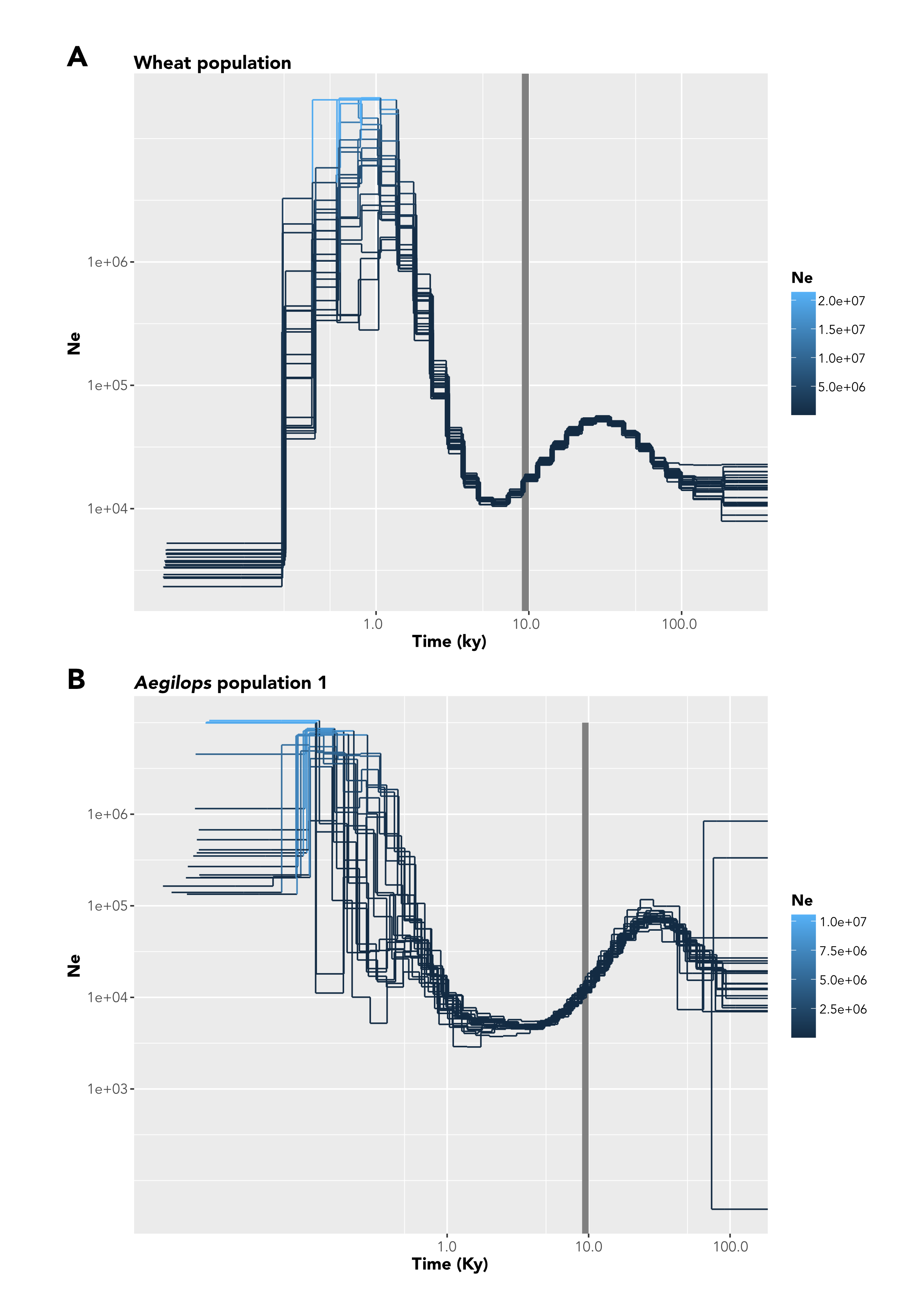

### S10_Fig.tiff

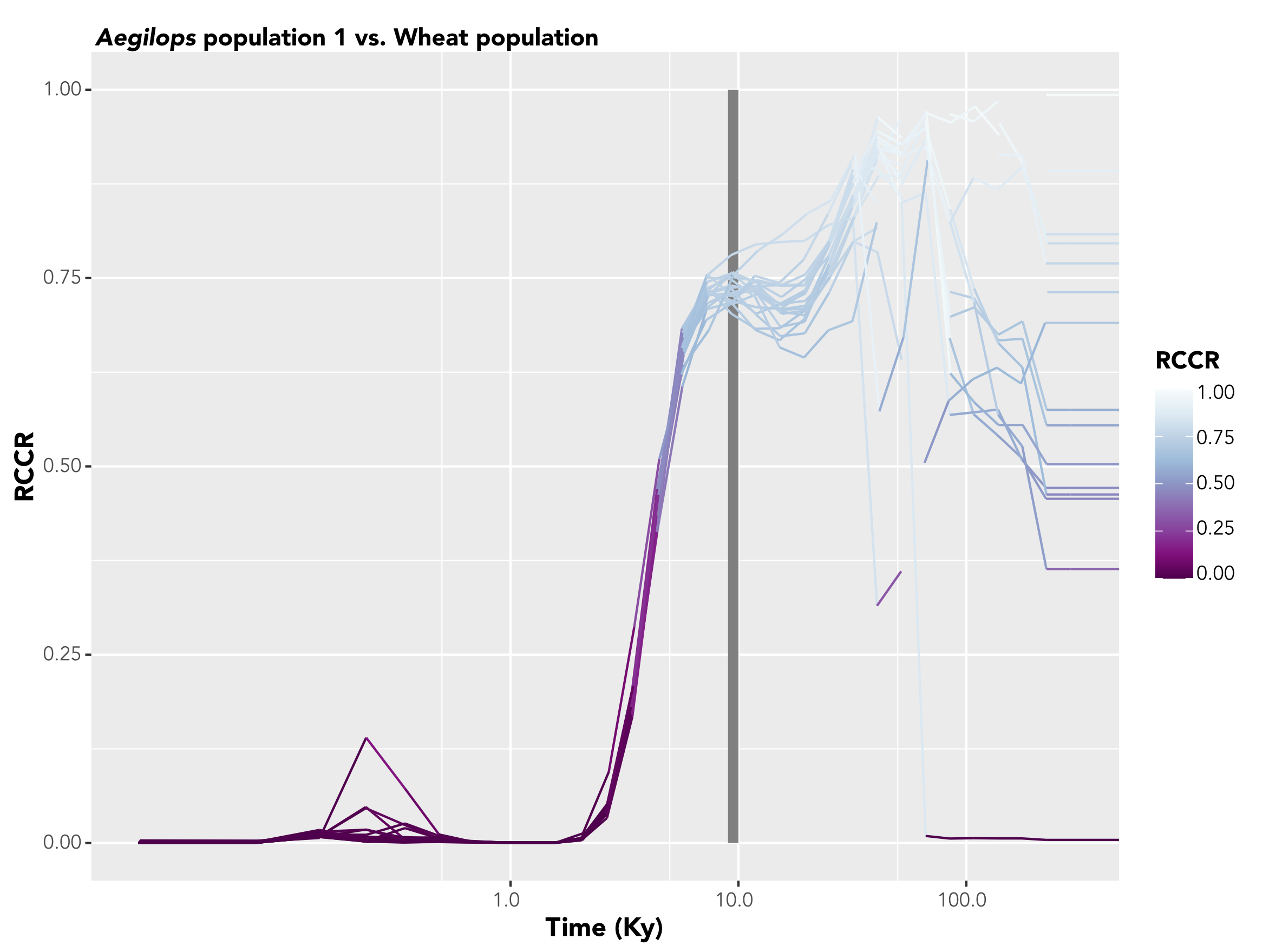

### S11_Fig.tiff

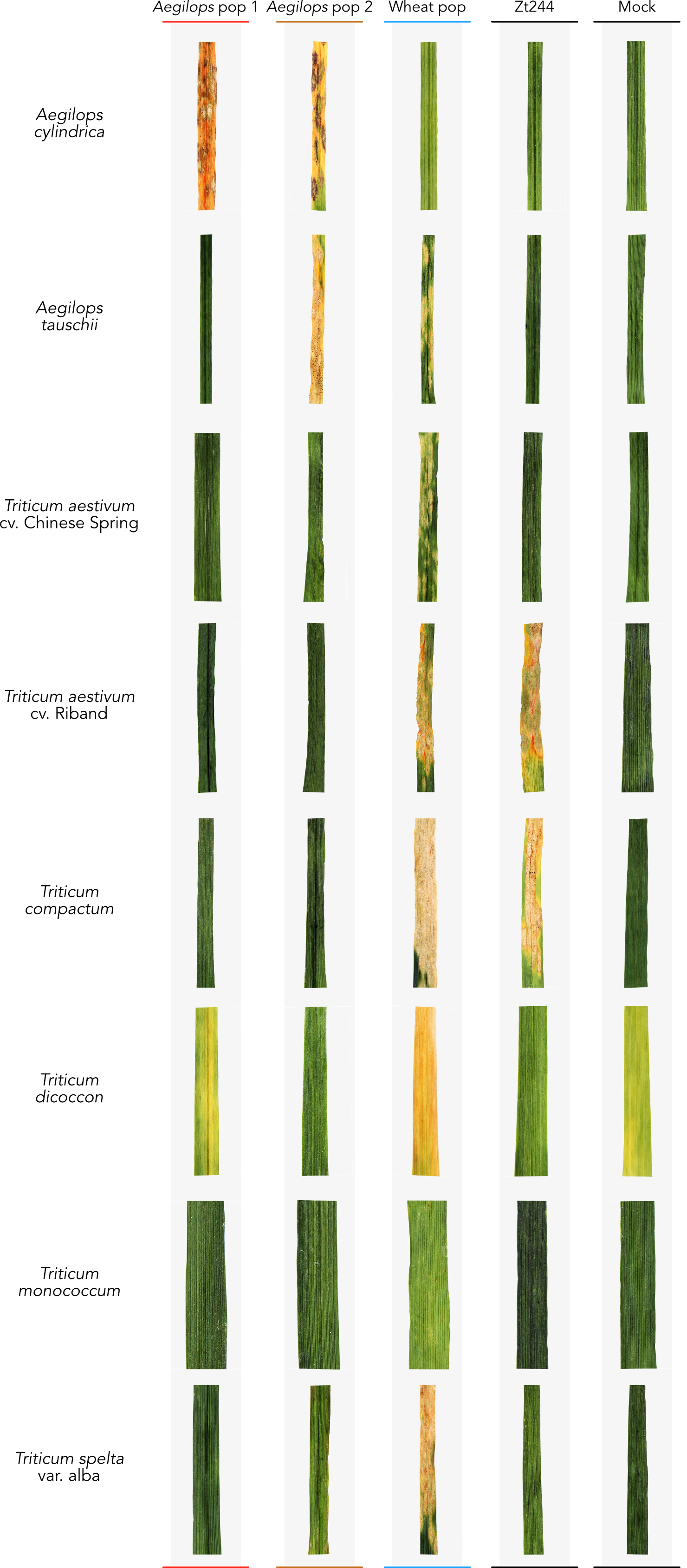

### S12_Fig.tiff

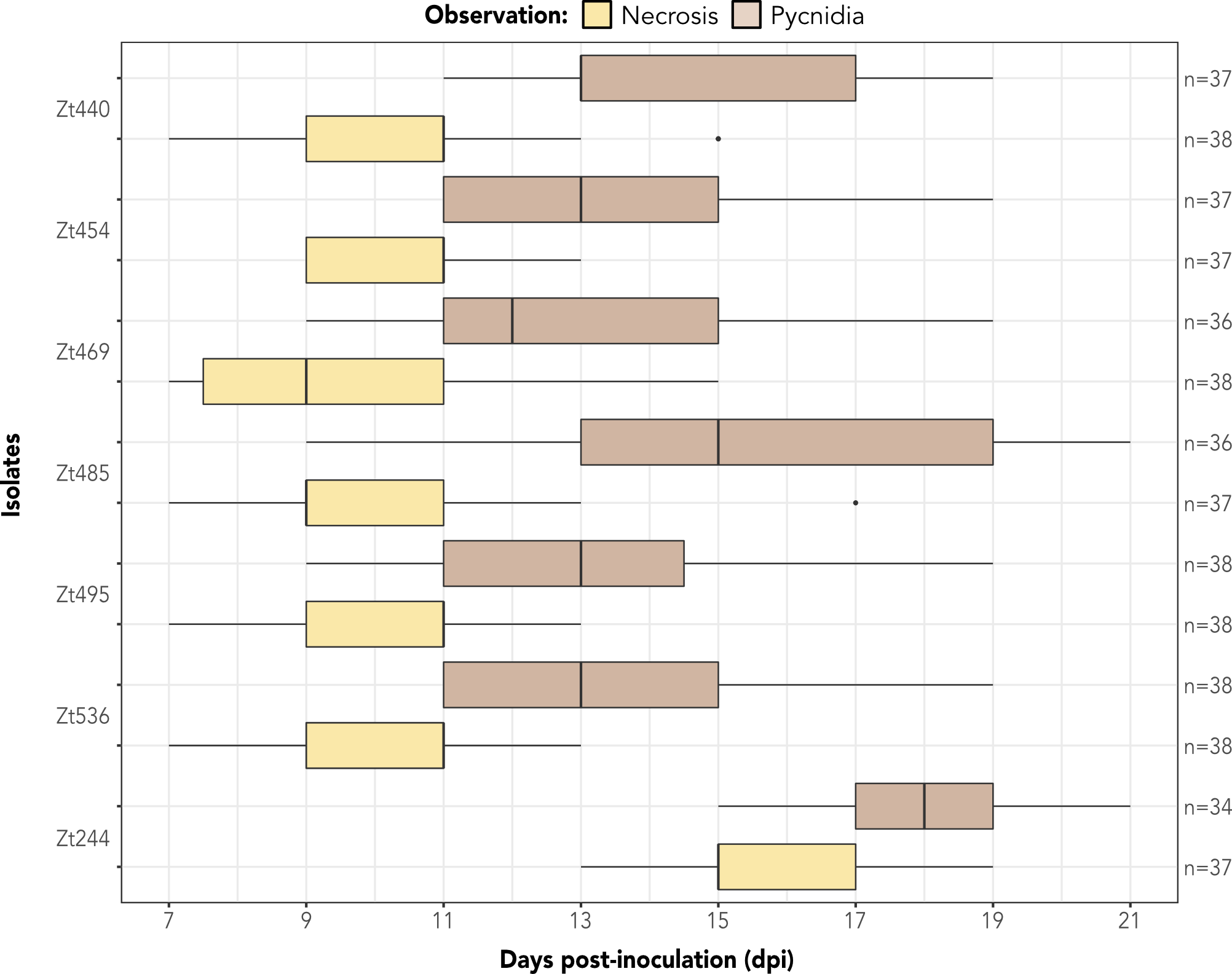

### S13_Fig.tiff

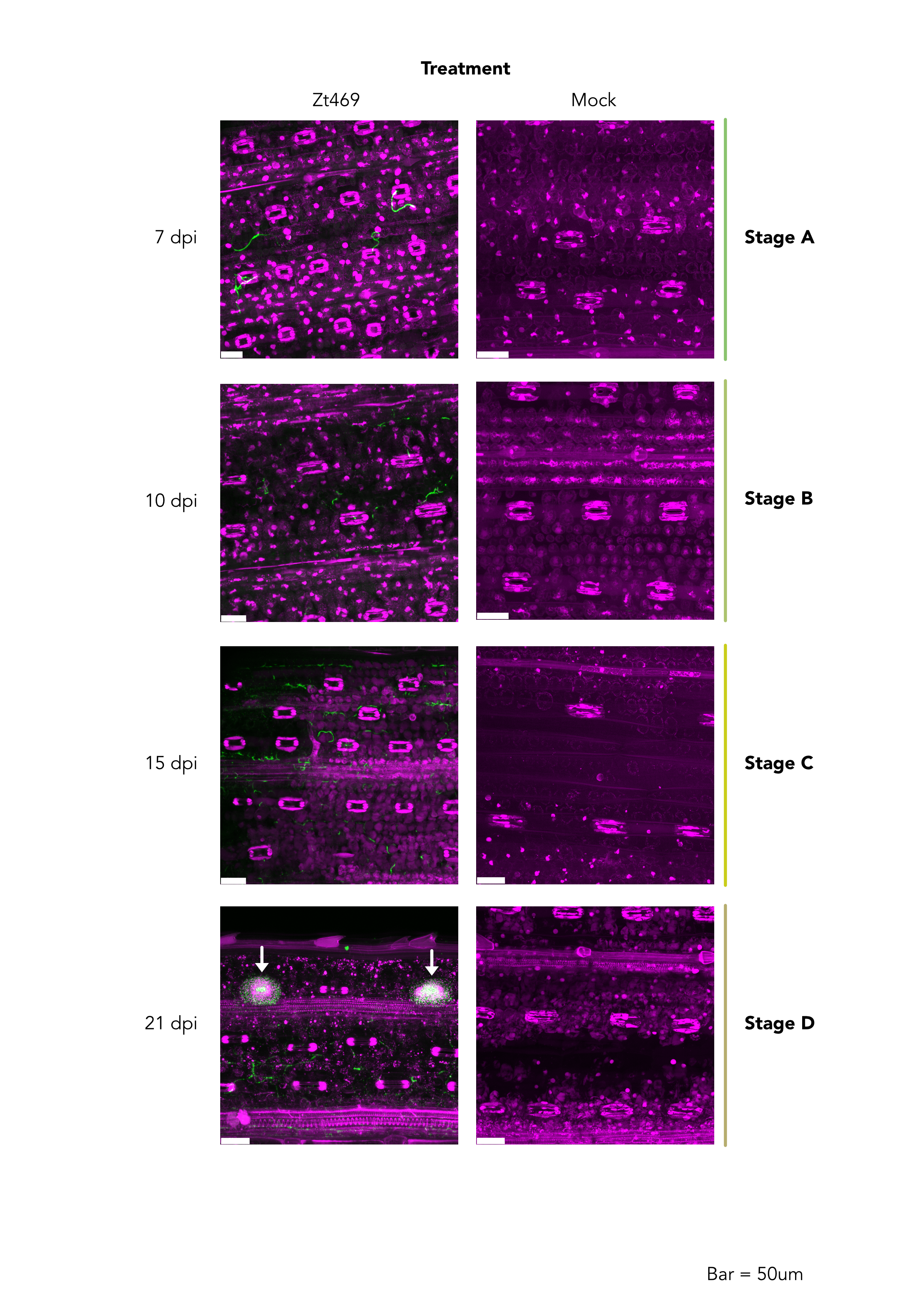

### S14_Fig.tiff

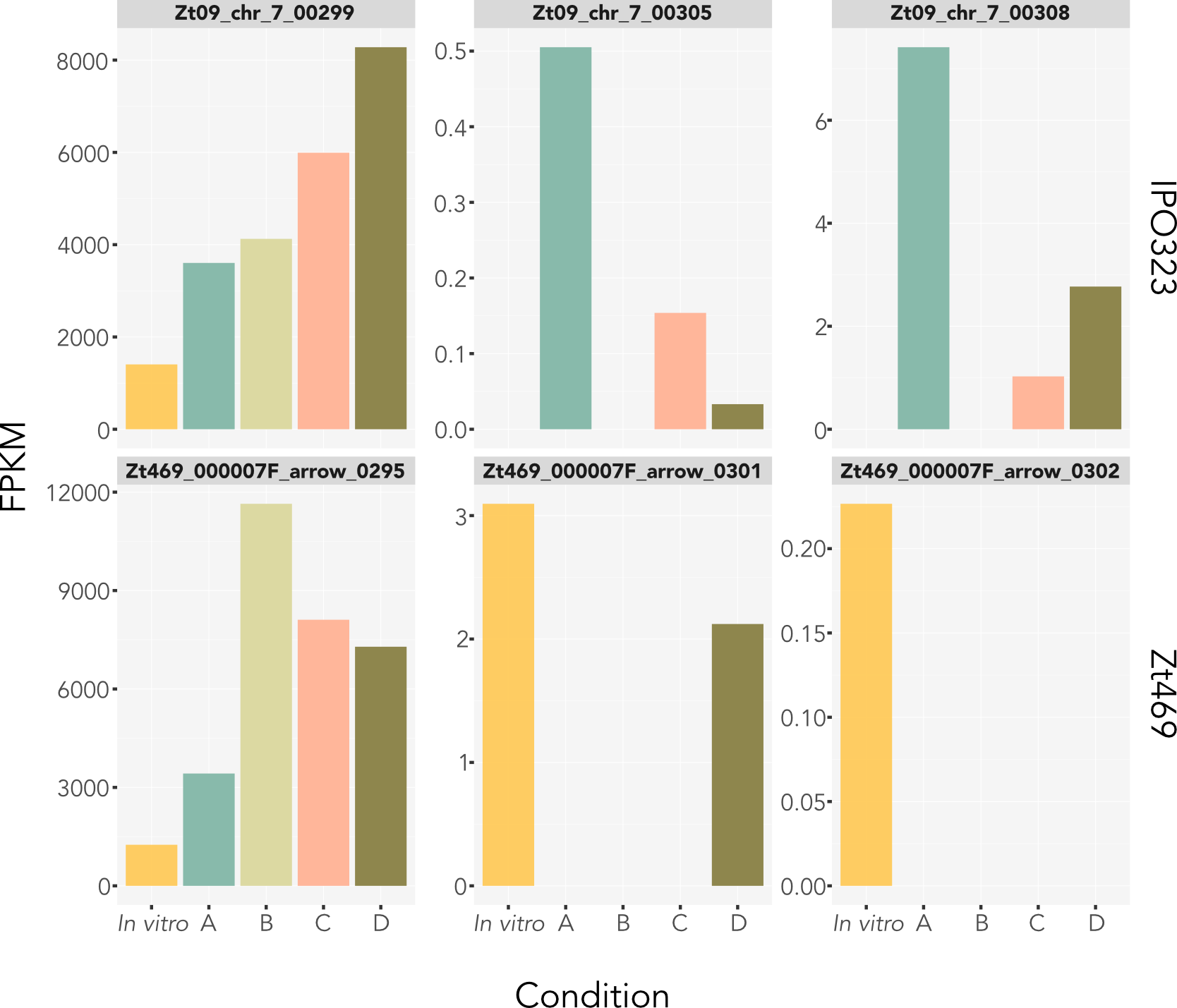
